# SIMO – Single Section Integrative Multi-Omics – spatial mapping of metabolites and lipids combined with region-specific proteomics in a single tissue slice

**DOI:** 10.64898/2026.04.17.719206

**Authors:** Kevin Hau, Antonia Fecke, Felix-Levin Hormann, Ann-Cathrin Groba, Luiza Martins Nascentes Melo, Feyza Cansiz, Gabriele Allies, Andreas Hentschel, Jianxu Chen, Sven Heiles, Alpaslan Tasdogan, Albert Sickmann, Karl Smith

## Abstract

Technological advances in biomedical sciences have accelerated multi-omics research, enabling high-resolution spatial mapping of diverse molecular compound classes. However, integrating spatial omics often requires serial tissue sections, limiting the alignment correlation across modalities. We present a single-section integrative multi-omics (SIMO) workflow that combines metabolite and lipid imaging with histopathology and region-specific proteomics. Using MALDI-MSI, tissue staining, and laser microdissection (LMD), SIMO delivers comprehensive metabolic, lipidomic, and proteomic insight from the same sample. Using mouse cardiac tissue we develop, control, and validate the methodology resulting in ∼60 imaged lipids and ∼60 imaged metabolites at 20 µm pixel size and subsequently spatial proteomics by LMD, detecting over 5,000 proteins from the same tissue. To demonstrate the capabilities of the workflow in preclinical context, we apply SIMO to a metastasizing melanoma PDX model, identifying over 100 spatially localized lipids and metabolites, and over 5,000 proteins across metastases and non-tumor tissues in liver. SIMO enables precise ROI selection, statistical comparison of protein regulation, and alignment of metabolic and lipidomics pathways across spatial omics and region-specific proteomics, demonstrating its value as a spatial multi-omics platform.

## Main

Advancement of analytical technologies and bioinformatics in recent years has revolutionized the possibilities of analysis in biomedical sciences, including the prospect of combining spatially-resolved untargeted metabolomics, lipidomics, and proteomics^1,2^. Multi-modal and spatially-resolved workflows hold the promise to provide a systems biology perspective on intertwined biological regulatory machineries in heterogenous diseases such as cancers that are difficult to unravel with classic omics workflows^3–5^. In this context, spatially resolved tissue organization has proven critical for developing therapeutic strategies, for example enabling prediction of immunotherapy response in non-small lung cancer (NSLC)^6^ and supporting systematic analyses of skin specimens in toxic epidermal necrolysis (TEM)^7^. Collectively, these studies underscore the growing centrality of spatial biology and its enabling technologies in contemporary biomedical investigation^8^.

One recent prominent example is deep visual proteomic (DVP) analysis in which fluorescent microscopy and mass spectrometry (MS) are combined ^7,9,10^. By utilizing laser microdissection (LMD) capture technology and pathological staining, identified cells are individually excised from the tissue and analyzed with proteomics workflow resulting in numbers of 1000-5000 proteins in a single cell or cluster of cells identified for example in liver tissue^11,12^ and colon organoids^13^.

While these case studies illustrate the biomedical promise of spatial technologies, integrating multiple omics layers from the same tissue section without compromising histopathological context remains a major challenge. This limitation is particularly consequential in clinical tissue analysis, where the concurrent attainment of high molecular sensitivity and high spatial resolution would enable direct alignment of metabolic, lipidomic and proteomic signatures with established morphological features. Such joint readouts would provide an integrated, mechanistic view of tissue biology and disease, opening new opportunities to resolve pathogenic processes and to inform targeted therapeutic strategies grounded in the underlying molecular architecture.

Matrix-assisted laser desorption/ionization mass spectrometry imaging (MALDI-MSI) is a widely used spatial omics modality and has been incorporated into diverse multi-omics workflows^14^. These efforts include transcriptomic pipelines, enabling single-cell–level transcript comparisons, as well as coupling to targeted derivatization strategies to enhance neurotransmitter detection^4^. A common strategy in MSI-based multi-omics strategy is to analyze serial tissue sections to profile distinct molecular layers within a shared anatomical frame. However, the reliance on serial sections substantially complicates spatial integration across features and different omics levels becomes challenging. This is because molecular information derived from the different omics layers do not originate from the same cellular layer. To mitigate this limitation, several studies have advanced workflows that extract multiple omics modalities from the same tissue section or sample, thereby improving cross-modal comparability and analytical consistency. Quanico *et al.* combined MSI with subsequent LC-MS/MS-based region-of-interest (ROI) analysis using grid-aided, parafilm-assisted microdissection based on manual scalpel excision of ROIs^15^. Mezger *et al*. established a single-section approach on conductive slides that coupled MALDI-MSI with direct proteomics, yielding ∼200 protein identifications in a specified ROI following MSI acquisition^16^. Similarly, Hendriks *et al.* reported a MALDI-MSI-LC-MS/MS workflow enabling quantitative lipidomic imaging alongside proteomic profiling from a single tissue section^17^.

Despite substantial progress towards integrated single-section multi-omics, key limitations remain with exploitations still available. Many existing workflows prioritize proteins, lipids, or other non-polar metabolites, whereas coverage of polar small mass metabolites, which play a central role in cellular energy metabolism and signaling, together with lipids and proteins remains limited. In addition, histopathological staining, such as hematoxylin and eosin (H&E) staining is frequently not incorporated constraining the alignment of spatial imaging readouts and molecular profiles with established morphological annotations. So far it has not yet been reported of a workflow that combines MALDI-MSI of small mass polar metabolites and lipids with histopathological staining and untargeted proteomics from a single tissue section mounted on a glass slide. This approach can be highly beneficial, allowing the combination of molecular mass imaging and histopathology to selectively guide the downstream selective protein area analysis.

Here, we demonstrate that SIMO enables seamless integration of histopathological H&E data with spatial lipidomics, metabolomics, and proteomics from one slide. The combination of MALDI-MSI and LMD as described here enables spatial co-registration without sacrificing molecular coverage facilitating a systems biology view of tissue samples. This is demonstrated for a melanoma patient-derived xenograft model for metastasis formation. Phenomenologically, SIMO captures the differentiation of molecular profiles between liver metastases of varying sizes. To demonstrate the capabilities of our workflow, SIMO is able to entangle that metabolic demand in small metastasis is dominated by processes that accelerate growth such as higher succinate dehydrogenase and fumarate levels compared to larger metastases in the same tissue.

## Results

### Single-section analysis enhances spatial information and increases data output from minimal sample input

We commenced with designing a multi-omics workflow that is capable combining spatially resolved metabolomics, lipidomics, and proteomics with histological workflows all on the same tissue section. For metabolomics and lipidomics we chose MALDI-MSI as sampling method, whereas proteomics and corresponding microscopic investigations were performed on a LMD setup. We hypothesized that by adjusting the energy per pulse of the MALDI laser, no or only minimal ablation of the tissue would permit subsequent staining for microscopic investigations followed by spatial proteomics. Therefore, the workflow shown in **Figure 1** should be capable to achieve the multi-modal imaging on a single tissue section.

**Figure 1:**
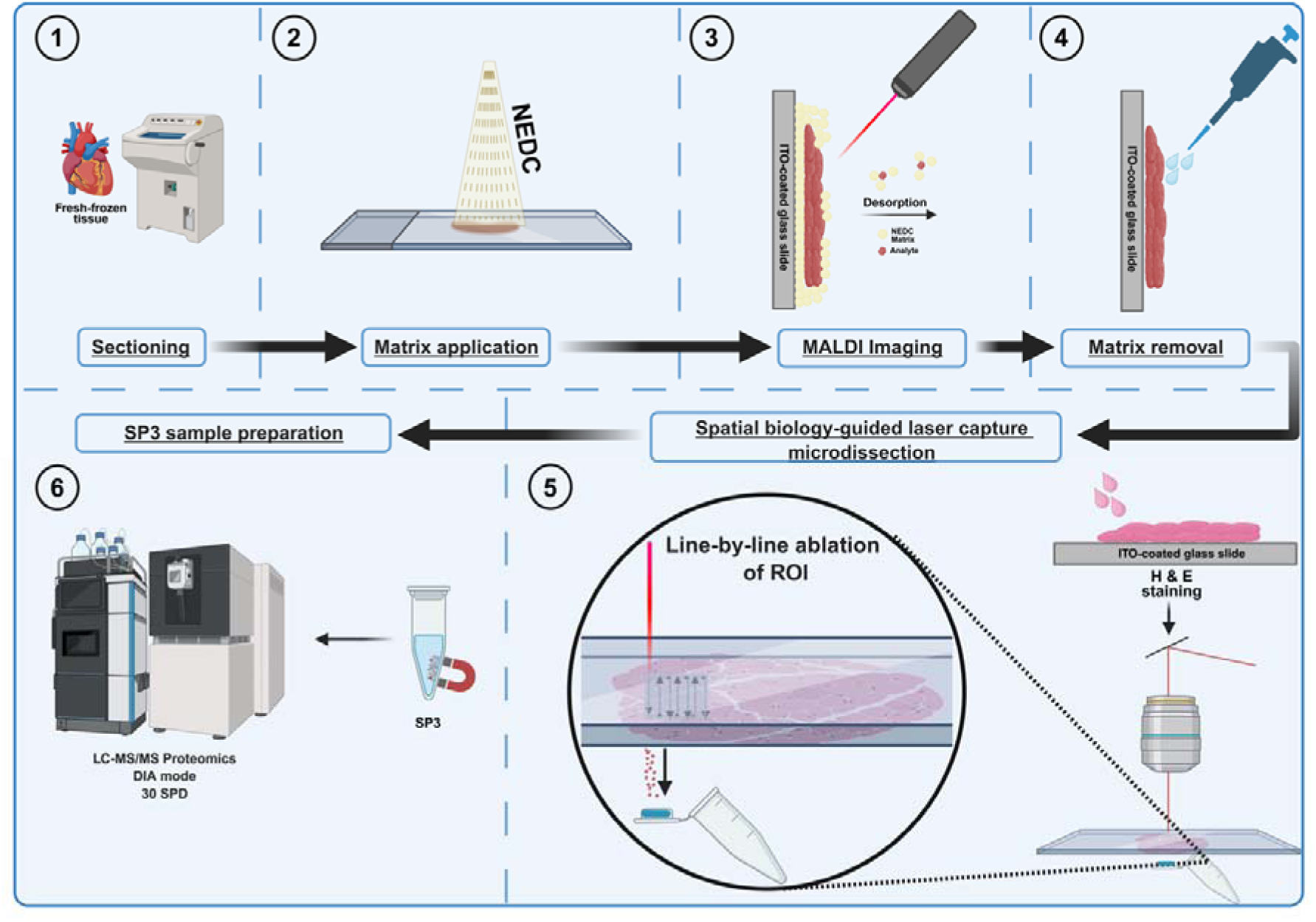
Workflow of the single-slide multiomics workflow (SIMO). (**1**) Snap-frozen tissue is cut into 12 µm thick sections at -20°C, placed onto ITO-coated glass slides and (**2**) NEDC matrix is applied onto the tissue. (**3**) MALDI imaging is carried out after crystallization of NEDC matrix. (**4**) After MALDI imaging, NEDC matrix is removed via EtOH and hematoxylin & eosin staining is performed before (**5**) Histopathology or MSI-guided laser capture microdissection. Since a regular glass slide coated with ITO is used, the “Draw-and-Scan” function of the LMD is applied to ablate regions of interest (ROIs) from upside-down positioned tissue line-by-line into the collection tube cap, filled with lysis buffer. (**6**) Protein sample preparation is executed by single-pot solid-phase sample preparation (SP3), injected into a Vanquish Neo UHPLC system coupled to an Orbitrap Astral mass spectrometer and analyzed in DIA mode with a 30 samples-per-day (SPD) method.

Next, we tested which experimental conditions enabled the combination of MALDI-MSI, spatial proteomics, and microscopy without sacrificing the performance of the individual modalities. For this purpose, various conditions and sampling schemes were compared as summarized in **Figure S1**. As the spatial proteomics pipeline is at the end of the envisioned workflow, we tested to what extent the H&E staining, washing steps, and MALDI measurements affect the protein numbers. Whereas the control heart tissue workflow gave the largest number of proteins, all sample preparation steps, i.e., washing and H&E staining, recover about 75 % of all proteins in the control tissue (**Figure 2A, Figure S2A**). Complete protein pathway coverages were not lost in the final workflow but show that major pathways such as central carbon metabolism are not affected (**Figure S2B**). When MALDI-MSI is additionally performed before proteomics, the overlap to the control is not further affected indicating that H&E staining is the major cause for loss of proteins. As revealed by the correlation analysis shown in **Figure 2B**, protein intensities exhibit correlation coefficients above 0.90, leading us to conclude that the sample preparation steps prior to our proteomics assay do not alter protein abundances but rather lead to the loss of some proteins during the H&E staining process. Next, we tested if the same number of lipids and metabolites are obtained when performing MALDI-MSI with a split or full *m*/*z*-range and how this affects protein numbers. Whereas protein numbers when employing the full *m*/*z*-range declined compared to two subsequent MALDI-MSI measurements due to the increase in ablated area compared to two measurements with a spatial offset and metabolite/lipid numbers did not significantly differ between the methods (**Figure S2-4**), the larger pixel size and the increased measurement time required for measurements with spatial offset outweighed the reduction in protein annotations. Therefore, we decided to employ MALDI-MSI with the full *m*/*z*-range to enable decreased pixel sizes.

**Figure 2:**
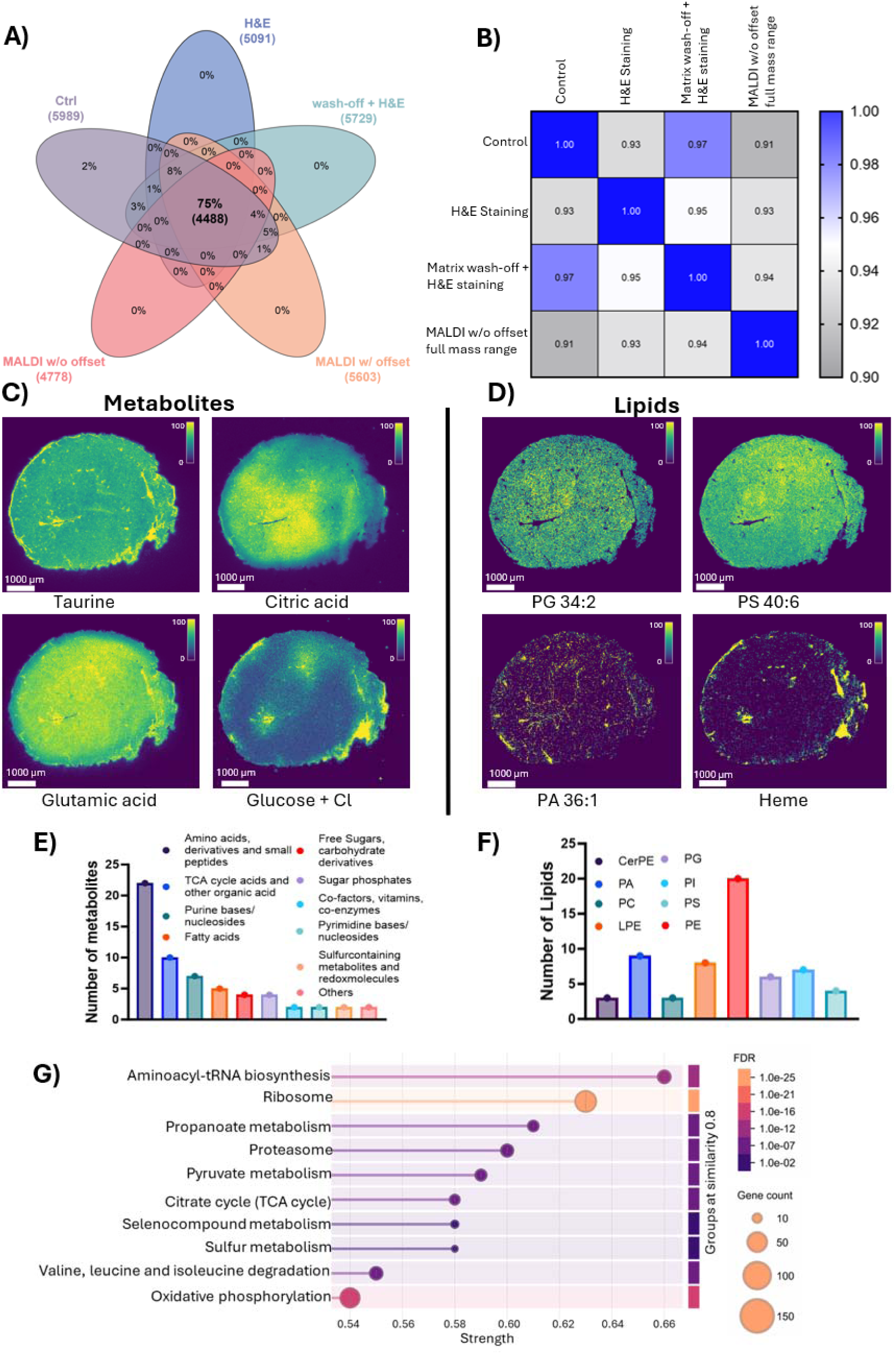
Method Development on mouse cardiac tissue. (A) Venn diagram visualizing the overlap of protein identifications at different stages of method development. Numbers and percentages represent shared and unique proteins in each step. (B) Pearsońs correlation coefficients of log2-transformed protein LFQ intensities across method development steps. Exemplary MALDI images of metabolites (C) and lipids (D) identified in heart tissue. Signal intensity is indicated by the color scale. (E) Chemical classification of identified metabolites in heart tissue in negative ion mode (F) Number of lipids from different classes identified in heart tissue in negative ion mode. G) KEGG Pathways enrichment analysis using STRING database (Version 12.0) based on the input protein set of the final applied method (MALDI w/o offset; 4778 proteins). Pathways are ranked according to the STRING strength score. The displayed pathway showed significant enrichment (FDR ≤ 0.05).

The results of the optimized SIMO workflow for mouse heart tissue are summarized in **Figure 2C-G**. The homogeneous heart tissue shows uniform distributions of most metabolites for which some examples are shown in **Figure 2C** and fine structures such as blood vessels are visible in the higher *m*/*z*-range (**Figure 2D**). As summarized in **Figure 2E-G**, the developed SIMO workflow covers the most important metabolites (**Figure 2E**), all typical lipid classes detected in negative-ion mode (**Figure 2F**), and a broad range of proteins (**Figure 2G**) with the ability to obtain histological characterization on the cellular level via H&E from the same section prior and after the SIMO method (**Figure S1,S4**).

### Multi-omic spatial regulation in a metastasizing melanoma PDX mouse model

We next applied the method in a preclinical setting. Specifically, we profiled a human patient-derived xenograph (PDX) model of melanoma liver metastasis^18^. Melanoma, a common malignancy originating from melanocytes (most often in the skin), is frequently curable at early stages; however, advanced disease is characterized by a high propensity for metastatic dissemination. Visceral metastases, including to the lung and liver, are associated with aggressive disease and poor prognosis. The herein employed human melanoma PDX model recapitulates key features of metastatic melanoma, producing multiple liver tumor nodules that vary in size and morphology. Macrometastatic lesions (> 5 mm) were readily identified by histopathological staining, enabling size-based stratification and systematic multi-omics analysis of metastatic progression (**Figure 3A**). The tissue section exhibited pronounced regional heterogeneity (**Figure 3B**). We captured this heterogeneity in the MALDI–MSI data using hierarchical spatial clustering with complementary UMAP embedding (**Figure 3C**), which revealed discrete molecular domains. The clustering resolved the major anatomical and pathological features without over-segmentation. The combined UMAP and hierarchical clustering framework, distinguished dominant tissue feature types while delineating gradual biochemical transitions without imposing hard boundaries and preserved nested structure within the data. Cluster inspection indicated a metabolic shift from larger macrometastases (cluster 1, light blue) toward micrometastases (< 5 mm) and the peripheral regions of larger tumors (cluster 3, peach), suggesting that proliferating tumor cells at macrometastatic borders share molecular profiles with smaller metastatic deposits. To illustrate the spatial localization, representative MALDI-MSI ion images for metabolites and lipids are shown in **Figure 3D&E,** respectively. Several energy-related metabolites displayed marked intensity differences in tumoral regions to adjacent non-tumor liver tissue (hereafter, referred to as non-tumor tissue healthy tissue). For example, phosphopantothenic acid, hexose monophosphate and HexCer 36:3, showed strong tumor–non-tumor contrast (3.3-, -2.1- and 4.7-fold change of averaged signal intensities, respectively; log2 fold change tumor/non-tumor), with detection signals largely absent in the opposing tissue state. By contrast glutamic acid and glutathione exhibited more modest differences (0.5, -0.5 log2 FC tumor/non-tumor), but varied across the spatial patterns, revealing gradients across regions of interest. PS 38:4 showed particularly heterogeneous spatial distributions, with generally reduced intensity in tumor regions but substantial variability across the section, potentially reflecting fine-scale liver architecture or zonation. Additional metabolite and lipid imaging examples and statistical analyses are provided in **Figures S5–S7**. Across three biological replicates, we tentatively annotated 98 metabolites and 93 lipids by MALDI–MSI, the majority of which were confirmed by LC–MS/MS (**Table S1**).

**Figure 3:**
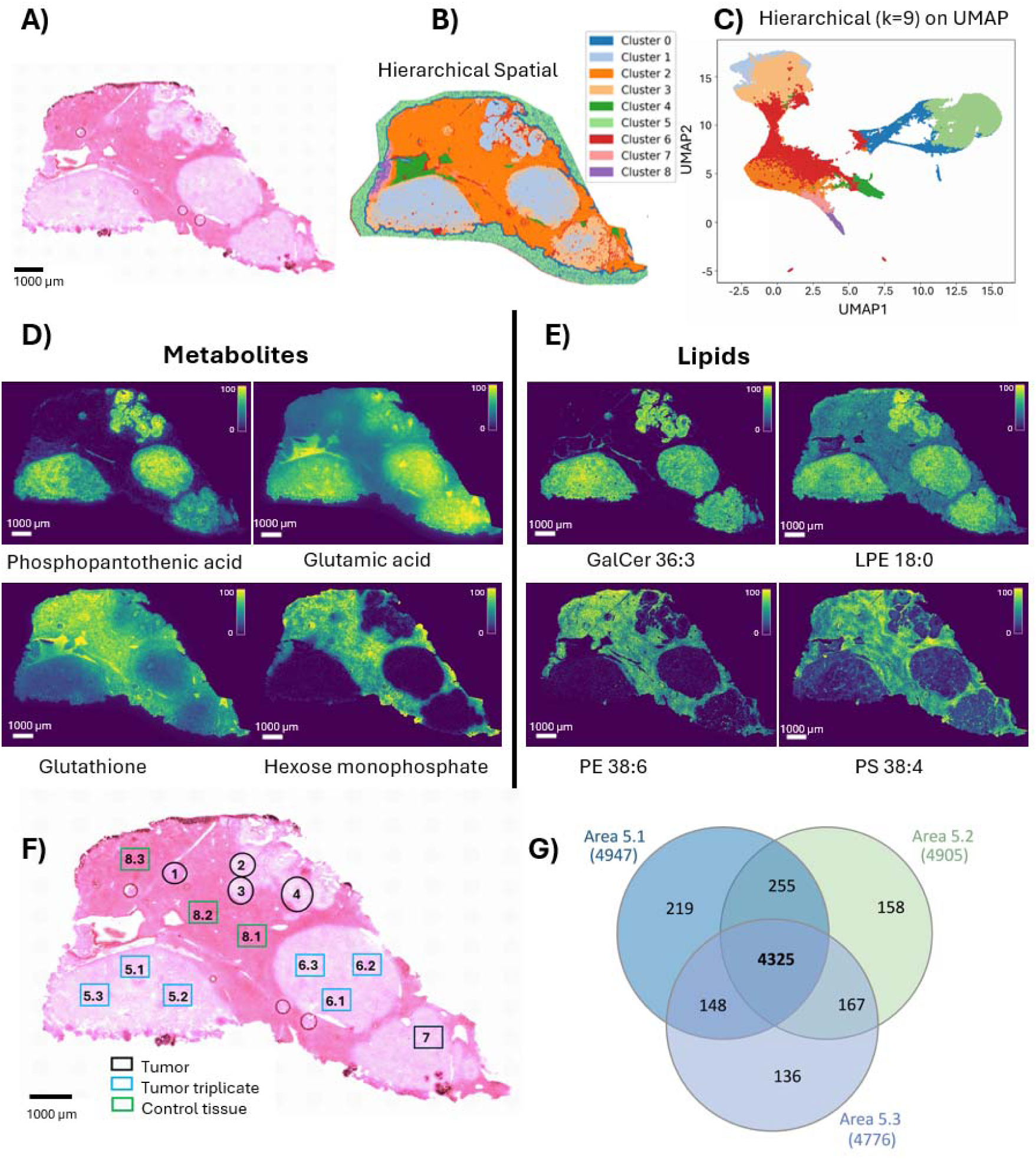
Application of SIMO to PDX melanoma mouse model. (A) H & E stained tissue section from the patient-derived melanoma metastatic liver (B) Hierarchial clustering image based on MALDI-MSI data with 9 clusters. C) Corresponding hierarchial UMAP showing clustering patterns between metastatic and non-tumor tissue. Exemplary MALDI images of metabolites (D) and lipids (E) with higher (top) and lower (bottom) intensity in melanoma liver metastases. Signal intensities are indicated by the color scale. (F) Depiction of LMD excised ROIs of different metastases and non-affected tissue. Metastases 1, 2, 3 and 4 were excised entirely, whereas for larger metastases, three randomly located replicates of 500,000 µm² were excised within the metastasis. (G) Venn diagram shows the overlap of identified proteins between three replicates of metastatic liver tissue area 5. For each replicate, 500,000 µm^2^ are excised during LMD.

**Figure 3F** shows the LMD-excised regions after MALDI-MSI and H&E staining, with areas numerically annotated for reference. ROIs were defined based using staining and MALDI-MSI data to delineate regional features, including tumor boundaries (**Figure S8**). Owing to their larger sizes, tumor tissues labelled 5 and 6 were sampled as three separate ROIs (5.1-5.3 and 6.1-6.3) of 500,000 µm^2^ each to assess intratumor heterogeneity (**Figure 3F**). Overlap in molecular protein features across the three sub-regions was evaluated and is shown in a Venn diagram (**Figure 3G**). We observed an overlap of 80% for each of the sub-regions 5.1 - 5.3, with over 4700 proteins identified in each region. This indicates that only minor intratumoral heterogeneity can be observed at the proteomics level for this larger regional size of analysis. By contrast regions 1, 2, 3, and 4 were excised in their entirety, because of their smaller size, enabling intratumoral comparisons. The feature comparison of intratumoral and intertumoral metastastic data highlights this method can deliver informative insights at high spatial region analysis, but shows potential for more in-depth intratumoral differences for larger regions of tissue that may differentiate upon advanced stages of growth. By interpreting the molecular data, more spatially defined ROIs could be analyzed for higher spatial regional analysis.

### Metastatic metabolic programs resolved across omics layers in a single tissue section

To evaluate the discriminative capacity of the SIMO method on a preclinical metastasis model, we performed a multi-omics comparison of non-tumor liver tissue and melanoma liver metastases (**Figure 4**). As mentioned earlier, lesions were stratified by size into small mircometastasis (small metastases, < 5 mm, tumor 1), intermediate-size micrometastasis (Intermediate metastases, 5-10 mm, tumor 2 – 4, clustered) and large macrometastasis (large metastases, > 10 mm, tumor 5 – 7). This size-based classification enabled a systematic assessment of stage-associated metabolic, lipidomic, and proteomic changes. Seamless MALDI analysis via H&E-stained histopathology guidance as implemented in SIMO enabled association of molecular signatures to metastatic and non-tumor tissue. **Figure 4A-B** show principal component analyses (PCA) of metabolite and lipid MALDI–MSI datasets comparing non-tumor tissue with metastatic lesions. In the metabolomics dataset, non-tumorregions were similar to small metastases, whereas intermediate and large lesions separated progressively into distinct clusters. By contrast, lipidomic profiles robustly distinguished non-tumor tissue from all metastases, with non-tumor tissue forming a clearly independent cluster. These differences were also evident in the spatial MSI data: lipid species showed sharply demarcated, lesion-restricted distributions, whereas small-molecule metabolites exhibited more gradual transitions between non-tumor and metastatic regions. Notably, this pattern was reversed when comparing intermediate and large metastases.

**Figure 4:**
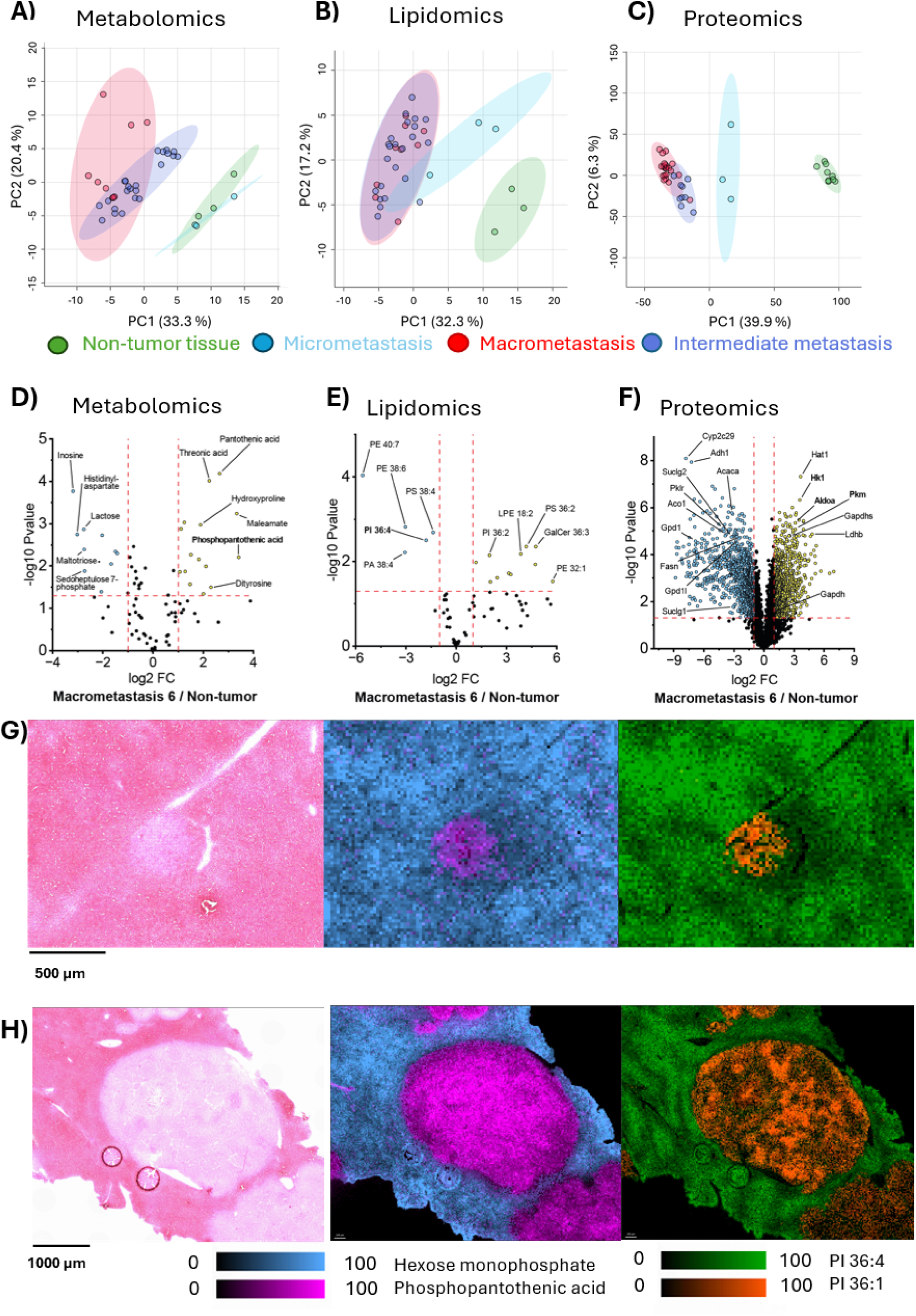
Comparative analysis of tumor and non-tumor tissue. Melanoma liver metastases were grouped as macrometastasis (areas 5, 6, 7 **Figure 3**), intermediate metastasis (areas 2,3,4 **Figure 3**) and micrometastasis tumor (area 1, **Figure 3**) and compared against adjacent non-tumor tissue (area 8) from all three sections. (A) Metabolomics based principal component analysis (PCA) plot from MALDI-MSI data after ROI analysis. (B) PCA plot generated from MALDI-MSI based lipidomics data after ROI analysis. (C) PCA plot generated from ROI based proteomics data after LMD excision. (D) Metabolomics-based Volcano plot comparing tumor area 6 (**Figure 3**) with adjacent non-tumor tissue from all three replicate sections. (E) Lipidomics-based Volcano plot comparing tumor area 6 with adjacent non-tumor tissue from all three replicate sections. (F) Proteomics-based volcano plot comparing tumor area 6 with adjacent non-tumor tissue from all three replicate sections. (G) H&E-stained tissue section zoomed in on tumor area 1 (**Figure 3**). Corresponding MALDI images show spatial distributions between non-tumor and micrometastatic areas of hexose monophosphate (blue) and phosphopantothenic acid (pink), and PI 36:4 (green) and PI 36:1 (orange) (H) H&E-stained tissue section zoomed in on tumore area 6 (**Figure 3**). Corresponding MALDI images show spatial distributions of two representative metabolites and two representative lipids overlayed in false colour. Corresponding MALDI images show spatial distributions between non-tumor and macrometastatic areas of hexose monophosphate (blue) and phosphopantothenic acid (pink), and PI 36:4 (green) and PI 36:1 (orange). Signal intensities are indicated by the color scale.

Small-molecule metabolites displayed increased heterogeneity and greater cluster separation, whereas lipid profiles from intermediate and large metastases overlapped extensively. Together, these results indicate that lipid alterations provide strong discrimination between non-tumor and metastatic tissue, while metabolic heterogeneity increases with lesion size and metastatic burden. **Figure 4C** shows analogous clustering for the spatially localized proteomics samples. Consistent with the MSI results, healthy tissue exhibited a distinct proteomic profile relative to tumor ROIs, and small metastases showed an intermediate profile between non-tumor tissue and larger lesions. At the protein level, the small tumor group separated clearly from intermediate/large tumors and non-tumor controls, a distinction not observed for lipids or metabolites. This suggests that early lesions may be characterized by proteomic remodeling that precedes or is not directly coupled to detectable shifts in lipid and metabolite abundance. Across omics layers, these trends capture spatially resolved biological heterogeneity and are consistent with the UMAP analysis of the same samples, indicating a gradual transition from small clustered micrometastases to larger metastatic lesions. Moreover, despite their closer size to the small lesions, intermediate tumors were molecularly more similar to large tumors than to small proliferative metastases.

To further characterize differences between non-tumor and diseased tissue, we performed a comparative analysis between tumor 6 and adjacent non-tumor regions using proteomics, metabolomics, and lipidomics. Differential abundance was assessed at each omics layer using volcano plots based on mean signals from non-tumor tissue and the tumor 6 cluster **(Figures 4D–F**). Of 98 detected metabolic features, 14 metabolites were significantly increased and 10 decreased in tumor tissue (**Figure 4D**). Among the increased metabolites were pantothenic and phosphopantothenic acid. Phosphopantothenic acid has been reported to accumulate in breast cancer, particularly in c-MYC high areas of human and murine tumors and serves as a precursor for coenzyme A biosynthesis, which supports TCA-cycle activity through intermediates including phosphopantothenic acid^19^. Consistent with these reports, our data suggest that altered pantothenate/coenzyme A metabolism may also contribute to metabolic adaptations in melanoma metastases. At the lipid level, mono- and di-saturated species (for example, PI 36:2 and PE 32:1) were enriched in tumor regions, whereas polyunsaturated fatty acid (PUFA)-containing lipids were reduced (**Figure 4E**). This pattern is consistent with elevated oxidative stress in metastases, which can favor increased lipid saturation. Proteomic profiling revealed multiple significantly altered proteins (**Figure 4F**), and we highlight candidates that align with the metabolic and lipidomic findings and were consistently detected and annotated in our datasets. Notably, hexokinase 1 (Hk1), aldolase A (Aldoa) and pyruvate kinase M (Pkm) were increased, consistent with enhanced glycolytic activity; these enzymes have been linked to tumorigenesis, invasion, migration, metastatic colonization and pro-metastatic programs in previous studies^20–24^.

We next illustrate spatially resolved molecular patterns in a small metastasis (**Figure 4G**) and a large metastatic lesion (**Figure 4H**). Overlaying MSI with histology revealed highly localized tumor-associated signals for hexose monophosphate and phosphopantothenic acid relative to adjacent non-tumor liver tissue. Lipid ion images showed analogous lesion-specific contrasts: PI 36:4 was predominantly detected in non-tumor tissue, whereas PI 36:1 was enriched in tumor regions, consistent with the reduced lipid unsaturation observed in **Figure 4E**. PI 36:1 and PI 36:4 differ only in their degree of unsaturation, highlighting lipid remodeling in metastatic tissue. More broadly, an increased ratio of monounsaturated (MUFA) to polyunsaturated (PUFA) fatty acids is a recurring feature of cancer metabolism and may reflect enhanced lipogenesis and/or protection from lipid peroxidation and ferroptosis through increased membrane saturation^25^.

In addition, the volcano plot in **Figure 4D** showed a significant decrease in sedoheptulose phosphate, a pentose phosphate pathway (PPP)–associated metabolite. Because the PPP branches from glycolysis and provides NADPH and nucleotide precursors, reduced sedoheptulose phosphate may reflect altered PPP flux in tumor tissue, with potential implications for redox homeostasis under elevated oxidative stress^26^.

### Stage-specific tumor differentiation by SIMO spatial multi-omics

Based on the SIMO data, different sized metastases exhibit distinguishable molecular profiles (**Figure 3B**). To further interrogate size-associated differences, we compared Tumor 1 (small micrometastasis) with Tumor 6 (large macrometastasis) across metabolomics, lipidomics and proteomics, with triplicate replicates performed. This analysis revealed significant size-dependent changes in metabolites, lipids, and proteins (**Figure 5A-C**). At the metabolite level,12 features were significantly upregulated in the large tumor, while 9 in the small tumor. Spatial zoom-ins of the corresponding ROIs recapitulated localized metabolic differences (**Figure 5D**). A prominent example was aconitic acid, which was strongly enriched in the large tumor but remained low in the small tumor and adjacent non-tumor tissue. As aconitic acid is a tricarboxylic acid (TCA) cycle intermediate, its accumulation may reflect altered TCA-cycle activity in larger lesions. One possible explanation is that macrometastases experience more pronounced hypoxia due to limited vascularization, promoting a shift toward anaerobic glycolysis and reduced TCA-cycle flux, with consequent accumulation of upstream intermediates such as aconitate. By contrast, smaller lesions may retain comparatively better oxygenation, supporting oxidative metabolism and sustained TCA-cycle activity. Proteomic differences were consistent with distinct metabolic states between lesion sizes. Several glycolysis- and PPP-associated proteins were elevated in the small tumor, including aldolaseB (Aldob), pyruvate carboxylase (Pc), pyruvate kinase (Pklr; liver isoform) and phosphoenolpyruvate carboxykinase 1 (Pck1), consistent with increased glycolytic and anaplerotic capacity relative to the large tumor. Aldob, has been implicated in routing carbon flux in liver metastases and supplying central pathways including glycolysis, gluconeogenesis and the PPP, and elevated Alodob expression has been associated with tumor progression^27,28^. Increased Pc may indicate enhanced metabolic flexibility by enabling anaplerotic replenishment of the TCA cycle under stress, supporting the notion that small lesions can more readily adapt to fluctuating nutrient and oxygen availability. We also observed increased Pck1 abundance in the small tumor, consistent with reports linking aberrant Pck1 expression to oncogenic signaling and metabolic reprogramming^29,30^. We additionally observed a non-significant size-associated trend in reduced glutathione (GSH). GSH is a key antioxidant, was lower in the large tumor than in non-tumor tissue and the small tumor, whereas the small tumor showed slightly higher signal than adjacent tissue. Glutathione has been implicated in metastatic colonization in other cancer types, and our data are compatible with a similar redox adaptation in newly established melanoma lesions. Consistently, glutathione S-transferase A3 (Gsta3) was strongly regulated in the small tumor (**Figure 5C**), supporting altered redox handling in this region. By comparison, lipidomic differences between small and large lesions were less pronounced, with three lipids increased in the large tumor and two increased in the small tumor. Nonetheless, we observed a trend in which more saturated lipid species were enriched in the small tumor, whereas smaller and more unsaturated species were relatively enriched in the large tumor. This was evident in the MSI data for LPE 18:2, which showed higher abundance in the large tumor compared with the small lesion.

**Figure 5:**
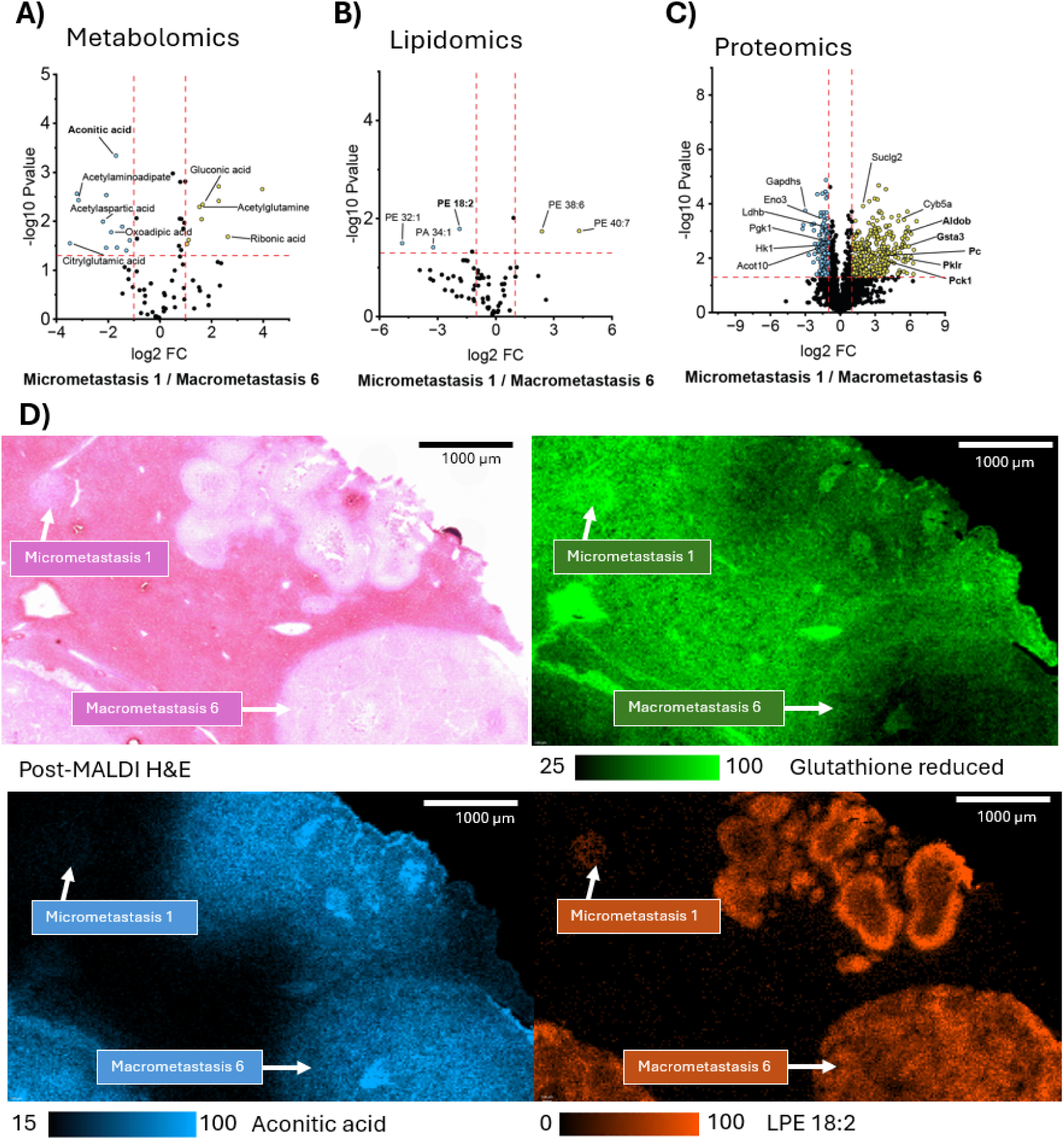
Analytical insight to metabolic differences of micrometastatic and macrometastatic tumors. (A) Metabolomics-based Volcano plot comparing metastasis area 6 (macrometastasis) with tumor area 1 (micrometastasis) from all three sections. (B) Lipidomics-based Volcano plot comparing micro- and macrometastasis from all three sections C) Proteomics-based Volcano plot comparing micro- and macrometastasis from all three sections. Annotation of tumor areas is consistent with **Figure 3**. (D) H&E-stained tissue section zoomed in on tumor area 1 and 6. Corresponding MALDI images show spatial distributions of selected representative metabolites and lipids that are differentially expressed in the different tumor regions. Glutathione reduced (green) is enriched in the small tumor, while aconitic acid (blue) and LPE18:2 (orange) are enriched in the larger tumor lesion. Signal intensities are indicated by the color scales.

Combining spatially integrated regions of metabolically different tissues with the regulating protein information enables pathway overviews to be considered. TCA metabolism is a highly anticipated pathway to study for clinical relevance, and with this advanced methodological usage, insights can be gained (**Figure S9**). Demonstrated in our method, a large coverage of TCA cycle metabolite imaging is achieved, with spatial differences and their corresponding protein regulations in micro- and macro-metastases. Higher regulation of succinate dehydrogenase (Sdh) in the micrometastatic area translates to the metabolic reaction of succinate to fumarate, with 2.4 times higher levels of Sdh detected in the micrometastatic region. Elevated levels of fumarate has been demonstrated to be a tumor growth promoter in the growth phase of tumor formation^31,32^. Using this protocol similar outputs can be visualized on a melanoma preclinical model, with spatially defined differences in a single tissue section of advanced growth tumors and proliferating ROIs.

## Discussion and Outlook

Integration and seamless combination of spatial analysis pipelines is a crucial step in elucidating the specifically localized mechanisms and biological functionality within life sciences. With SIMO we demonstrate a reproducible and robust workflow for single-section spatial lipidomics and metabolomics, histopathological staining and regionally defined proteomics analysis. We have established a multi-omic pipeline for MS-based analysis, by combining MALDI-MSI and precise excision of ROI tissue for proteomics by LMD. Method validation demonstrated the justification of using such a method, enabling high precision data with minimal variation across similar samples but displaying the high potential of heterogenous spatial analysis. By combining these, histological context can now be associated and explored in the specific area of interest for molecular profiling.

Previously reported multi-omic workflows demonstrated the inherent limitations, mainly the compromise of parameters to achieve multiple analysis types. For many, it is the loss of spatial depth and alignment, due to the necessity of serial sections for correlating methods. Others use further instrumentation features to enable multi-analyte class analysis, but there is usually a concession required, as demonstrated in **Figure S10**. Serial sectioning requires further depth off tissue to be used and can cause structural changes complicating the correlation of all data types with each other.

The introduction of each new step into our workflow was carefully assessed and compared against a control sample, demonstrating good recovery of protein identifications even after MALDI analysis. Application of our workflow to cancerous liver tissue with different sized metastatic lesions demonstrated consistent molecular separation between grouped tumors and adjacent non-tumor tissue at different omics levels. Our method captured biologically meaningful spatial patterns, such as region-specific metabolite and lipid distributions between tumor and non-tumor tissue from MALDI-MSI.

Additionally, SIMO revealed that lesions of varying size in the same patient-derived melanoma metastases model in liver tissue exhibit differentially regulated multi-omic processes. Whereas proteins associated with ß-oxidation (Acot10) and NADH/NAD+ metabolism (Ldhb) are upregulated in large metastases, smaller lesions have increased levels of enzymes and metabolites from the TCA (Sdh and fumarate) as well as compounds associated with countering oxidative stress (GSH and Gsta3). These results indicate that smaller lesions are armed with a differentially regulated repertoire of compounds to facilitate fast growth and negate damage to tumor cells. In contrast larger metastases in comparison switch to a “standard” tumor metabolism to sustain growth.

While our workflow demonstrates an insightful and explorative experimental framework for multi-omic data generation, further development dedicated to bioinformatic tools will be essential to fully integrate data from different omics levels. Improved computational frameworks will enable deeper interpretations and comprehensive biological insights. Beyond this proof-of-concept application, the workflow can be extended to multiple tissue types or can be further integrated with other omics layers, to gain deeper insights into cellular processes associated with tumor progression and other diseases.

## Methods

### Materials

MALDI matrix N-(1-naphthyl) ethylenediamine dihydrochloride (NEDC) was obtained from Sigma-Aldrich (Darmstadt, Germany). LC-MS grade solvents acetonitrile (ACN) and methanol (MeOH) were purchased from Biosolve (Valkenswaard, The Netherlands). Ethanol, Mayer’s hematoxylin, Xylene, Eosin Y (0.5% in alcoholic solution) and Histomount were all obtained from Sigma-Aldrich (Darmstadt, Germany). HPLC-grade water was purchased from Carl Roth (Karlsruhe, Germany).

### Animal model

In this study, heart organ tissue from a female Tie2-Cre mouse was used. Mouse breeding, housing, and organ collection were performed according to the guidelines from Directive 2010/63/EU of the European Parliament on the protection of animals used for scientific purposes. Organ removal for scientific purposes was performed according to § 4 of the Animal Welfare Act (Tierschutzgesetz; TierSchG). All animals were recorded, documented, and reported under the German Decree on the Reporting of Laboratory Animals (VersTierMeldV). A constant temperature of 22°C ± 2°C and a defined relative humidity of 55 ± 10% are ensured. The light/dark cycle is 12:12. Food and water are offered ad libitum. Animal sacrifice was done by cervical dislocation at an age of 12 months. Extracted organs were snap-frozen in liquid nitrogen and stored at −80 °C until analysis.

For the metastasis liver tissue, mouse experiments were conducted in accordance with relevant ethical regulations and approved by the local ethics committee in compliance with the European Convention for the Protection of Vertebrate Animals Used for Experimental and other Scientific Purposes (Directive 2010/63/EU, 81-02.04.2023.A026 Landesamt für Verbraucherschutz und Ehrnährung Nordrhein-Westfalen, Germany). For the melanoma metastasis model, melanoma cell suspensions were prepared for injection in Dulbecco’s Modified Eagle Medium (DMEM) medium supplemented with 1% penicillin/streptomycin, 10% fetal bovine serum (FBS) with 25% Matrigel (Corning). Subcutaneous injections were performed in the right flank of NOD.CB17*-Prkdc^scid^*/NCrCrl (NSG) (RRID:IMSR_CRL:394, Charles River, Germany) mice in a final volume of 100 µl. 6-weeks old female NSG mice were transplanted with 100 M481 melanoma cells^18^. Throughout the study, mice were maintained on standard chow and provided food and water ad libitum. The maximum allowable tumor diameter of 2.0 cm was strictly observed and not exceeded in any experiment. At that maximal ethical endpoint, all mice in the cohort were euthanized, per approved protocol, for analysis of primary tumor and metastatic disease burden.

### Tissue sectioning

Organ samples were cryosectioned prior to analysis using a Leica CM1860 cryostat. The tissue was placed into the cryostat chamber for 30 min of equilibration to the localized temperature. The organs were mounted to the cryosectioning chucks using deionized water and sectioned at 12 µm of thickness at -20°C. Sections were thaw mounted to ITO slides (70-100 ohms, Delta Technologies, USA). Sections were vacuum-packed and stored at - 80°C until further analysis.

### Matrix application

NEDC (7 mg/mL in 70:25:5 MeOH:ACN:H2O) was used as a MALDI matrix for negative ion mode and applied using the SunCollect Sunchrom sprayer. Heart samples were sprayed with 20 layers of matrix (layer 1-8: 15 µL/min, layer 9-20: 20 µL/min) at a speed of 800 mm/min and a nozzle height of z= 35mm. Metastasis samples were sprayed with 15 layers of NEDC (layer 1-3: 15 µL/min, layer 4-15: 20 µL/min) at a speed of 800 mm/min and a nozzle height of z=35 mm. Gas pressure was maintained at 3 bar.

### MALDI-MSI analysis

MALDI-MSI analysis was performed on an Orbitrap Q-Exactive mass spectrometer (Thermo Fisher Scientific GmbH, Bremen, Germany) coupled to an elevated pressure MALDI ion source (Spectroglyph LLC, Kennewick, WA, USA). A 349 nm MALDI Nd:YLF laser (Explorer One, Spectra Physics, Mountain View, CA, USA) was used and operated at a repetition rate of 500 Hz and pulse energy of 1-2 µJ. The laser was focused to a spot size of roughly 15 µm. Pixel sizes were varied in experiments. Heart images were acquired at 20x20 µm for full range analysis or measured two times at 30x30 µm with a 15x15 µm offset in the second analysis for separate metabolite and lipid measurements. Metastasis samples were measured at 20x20 µm. The MALDI source is operated at ∼ 6.5 Torr. The high-pressure funnel (HPF) and low-pressure funnel (LPF) were operated at 95 Vpp and 74 Vpp respectively using a 15% RF drive for 720 kHz and 825 kHz. The mass spectrometer was operated in negative ion mode with AGC set to fixed and a resolution of 120,000 at *m/z* 200. A set inject time of 250 ms was used, resulting in 125 shots/pixel. Scan range was varied in development experiments. Full-range experiments were performed at *m/z* 100-1200, separate metabolite and lipid analysis was performed at *m/z* 50-600 for metabolites and 600-1200 for lipids. All metastasis samples were measured from *m/z* 73.4 to 1100 *m/z*. Experiments were performed in triplicates.

### MALDI-MSI data processing

Generated Thermo RAW (.raw) and positional files (.xml) were converted to imzML format using the built-in-converter for LipostarMSI, which utilized msconvert from ProteoWizard (3.2.22317). The imzML files were loaded into Lipostar MSI using a 3ppm window, minimum peak intensity of 1%, peak detection frequency 1% and minimum spatial chaos 0.7. Data was TIC normalized at hotspot removal was applied using the 99% quantile. ROI analysis was performed using the same settings with no hotspot removal applied. **Figure S8** illustrates and highlights the manually drawn areas from the LipostarMSI software. From this, averaged spectra were generated and based on annotated compounds, statistical and visual differentials were analyzed.

Ion distribution maps are annotated based on MS1 data with a mass error below 3ppm. Annotation was performed based on an in-house standard library for metabolites^33^. PCA plots were generated in MetaboAnalyst’s one-factor statistical analysis. Metabolomics MALDI data is displayed with autoscaling and lipidomics data with auto-scaling and log10 transformation.

Lipid identification was performed in LipostarMSI based on the LIPDI MAPS database (https://lipidmaps.org/). After background subtraction, lipids were identified based on exact mass with a *m/z* tolerance of 5.0 ppm and cross validated with LC-MS/MS identifications of adjacent tissue sections. Results are manually checked for possible miss annotations based on known isomeric structures and isobaric overlaps.

MSI data were analyzed using custom Python scripts. The measured MSI data were imported from the paired .imzML and .ibd files using the *pyimzML* package. All spectra were binned along the *m/z* axis and normalized by the total ion current (TIC). To suppress residual noise, signals below a low-intensity threshold were set to zero. Log-transformation was applied to stabilize intensity variance prior to multivariate analysis.

For dimensionality reduction and visualization, principal component analysis (PCA) followed by uniform manifold approximation and projection (UMAP) was performed. or unsupervised clustering, hierarchical clustering was performed using the Agglomerative Clustering method from the *scikit-learn* package with ward linkage, which minimizes within-cluster variance. The clustering was applied to the PCA-reduced data, and the resulting segmentation maps were projected onto both the UMAP embedding and the original spatial image grid. A dendrogram was plotted with the dendrogram function from the *SciPy* package. For quality control Pearson correlation coefficients between replicates were calculated using the corrcoef function from the NumPy package (Python), based on the mean spectra.

### H&E staining

After MALDI-MSI analysis, sections were stained with hematoxylin and eosin (H&E). The MALDI matrix was first washed of for 30s in 95% EtOH. Heart samples were fixated for 3 min in PBS, immersed in diluted hematoxylin (1:2 dilution in 100% EtOH) for 45s. Then bluing was performed under running tap water for 5 min. Sections were counterstained in eosin Y for 1 min followed by a series of 15s ddH_2_O, 15s 70% EtOH, 15s 80% EtOH, 15s 90% EtOH, 3 min 100% EtOH and Xylene for 3 min.

Metastasis samples were stained following the same protocol. To increase contrast between regions, these samples were stained in hematoxylin for 1 min, followed by bluing for 1 min and counterstaining with Eosin Y for 2 min.

### Tissue Preparation for LC-MS analysis

Samples for LC-MS bulk analysis were collected while sectioning in the cryotome. For two phase extraction of metabolites and lipids, samples were homogenized in 150 µL ice cold ammonium acetate (0.1% in H_2_O) using the Precellys Evolution Touch homogenizer in three cycles (10s homogenization, 20s break, 5 min between each cycle). Next, 175 µL of MeOH (3% acetic acid) and 625 µL of MTBE were added and vortexed (1 hr, 4°C, 1200 rpm) for incubation. For layer separation, 200 µL ammonium acetate (0.1% in water) was added and samples were vortexed (15 min, 4°C, 1200 rpm) and centrifuged (19 min, 4°C, 10,000 rpm). The upper layer was collected and dried under nitrogen for lipidomics analysis. The lower layer was centrifuged again, transferred and dried. Dried samples were stored at -80°C until further processing.

Metabolomics samples were reconstituted in 80 µL (70:30 ACN:H_2_O), vortexed for 15 min (4°C, 1400 rpm) and incubated at -80°C for two hours. Samples were centrifuged (45 min, 4°C, 12,000 rpm) and transferred to fresh 2 mL Eppendorf. Prior to LC-MS/MS analysis, extracts were centrifuged again and transferred into LC-MS vials.

### LC-MS/MS methodology and data analysis for metabolite bulk tissue analysis

Metabolomics measurements were performed using a Vanquish Duo UHPLC system (Thermo Fisher Scientific, Waltham, MA; USA) equipped with a dual pump, autosampler, thermostatic column oven and a Atlantis Premier BEH Z-HILIC column. Mobile phase A consisted of 100% water with 10mM ammonium acetate and 0.1% ammonium hydroxide (pH 8.5) and 100% ACN was used as mobile phase B. Elution gradient conditions were as follows: 95% B, 0–2 min; 95–80% B, 2–7.7 min; 80–70% B, 7.7–9.5 min; 70–10% B, 9.5–10.5 min; 10% B, 10.5–12 min; 10–30% B 12–16 min; 30-95% B, 16-16.5 min; 95% B, 16.5-19 min. In this study an Orbitrap Fusion Lumos Tribrid Mass Spectrometer (Thermo Fisher Scientific, Waltham, MA, USA) with a heated electrospray ionization source (HESI) was used. Data was acquired in positive and negative mode separately at a resolution of 120,000 within the mass range 65-850 *m/z*. The HESI source was operated at a voltage of 3000V for negative mode and 3500V for positive mode analysis. Source parameters such as sheath, auxiliary and sweep gas were set to 50, 15 and 1 arbs respectively. The ion transfer tube was heated to 330°C and the vaporizer temperature was set to 350°C. Annotation was performed using an in-house spectral library based on MS1 detection and retention time matching. LC-MS/MS data was analyzed using Progenesis QI (v. 3.1) with an in-house standard library.

### LC-MS/MS methodology and data analysis for lipid bulk tissue analysis

The dried organic phases were reconstituted in 100 µL MeOH/i-PrOH (isopropanol) (1:1, v/v) and diluted by a factor of 10 to 100 depending on the sample. Samples were analyzed using an UHPLC system (Vanquish Flex, Thermo Fisher Scientific, Germany) equipped with a Triart C18 reversed-phase column (150 x 2.1 mm: 1.9 µm, 120 Å, YMC, Japan) coupled to a HESI-orbital trapping mass spectrometer (Exploris 240, Thermo Fisher Scientific, Germany). Separation was achieved by a 38 min gradient of mobile phase A (MPA) (ACN/H_2_O, 1:1, v/v) and mobile phase B (MPB, i-PrOH/ACN/H_2_O, 85:10:5, v/v/v) both containing 5 mM NH4HCO2 and 0.1 % FA at a flow rate of 300 µL/min, a column oven temperature of 50 °C, and an injection volume of 5 µL or 10 µL for positive or negative ion mode measurements, respectively. Gradient: 0 – 3 min from 10 – 25 % MPB, 3 – 9 min from 25 – 54 % MPB, 9 – 14 min from 54 – 77 % MPB, 14 – 18 min from 77 – 88 % MPB, 18 – 22 min from 88 – 95 % MPB, 22 – 30 min isocratic flow, 30 – 30.5 min from 95 – 10 % MPB, 30.5 – 38 min isocratic flow. MS measurements were carried out in data dependent top 10 acquisition mode. Survey scans were performed at a resolution of 180000 in a scan range of *m/z* 150 – 1200. For dd-MS2 measurements a resolution of 15000, minimum signal threshold of 5000, isolation window of 1.5 *m/z*, and stepped HCD activation energies of NCE 20, 25, 30 were set. Spectra were analyzed using Lipostar2 (v 2.1.4., Molecular Discovery, Italy). Data sets were loaded with a signal threshold of 1000. Blank subtraction, retention time filtering and removal of lipids without MS2 and isotopic pattern was performed prior identification. Identification is based on combined positive and negative ion mode measurements using *m/z* tolerances of 2.0 ppm and 6.0 ppm for MS1 and MS2 respectively.

### LMD protocol

The annotation of the metastatic tissue was performed by trained pathologists. According to the annotation, the regions of interest (ROI) of the H&E-stained tissue section were excised using laser microdissection (Leica LMD6 system, software version 7.6). In all cryosections utilized, metastases, designated with the numerals 1-4, were meticulously excised, whereas in the indexed areas 5.1 – 8.3, 500,000 µm^2^ were excised in each instance. The dissection was performed in brightfield mode with the following laser settings: The power is set at 60, the aperture at 5, the speed at 30, and the line spacing at 8. As the slides utilized are not of the membrane-based variety (such as PET or PPS membranes) but rather commercially available indium tin oxide (ITO) coated slides, it is not possible to cut out the tissue using the ‘Draw and Cut’ mode. Instead, the tissue must be ablated from the slide using the ‘Draw and Scan’ mode. In order to execute this procedure, it is necessary to position the glass slide in an inverted orientation within the sample holder. The 0.5 mL LowBind tubes were then positioned within the collection tube holder, followed by the addition of 40 µL of lysis buffer. The lysis buffer comprised 1% sodium dodecyl sulphate (SDS), 10 mM ammonium bicarbonate (ABC), 5 mM Tris-HCL, 15 mM sodium chloride, 1x protease inhibitor, and 1x PhosSTOP. Optical images of the stained tissues post-LMD are displayed in **Figure S11**.

### SP3 sample preparation

Proteolytic digestion was performed using single-pot, solid-phase, sample preparation (SP3^34^). First, the tissue was lysed by ultrasonic treatment (Bioruptor) in 10 cycles (1 cycle = 30 seconds of ultrasound, 30 seconds of cooling). To increase the proteolytic yield, disulphide bridges between cysteine amino acids were reduced with a final concentration of tris(2-carboxyethyl)phosphine (TCEP, Sigma Aldrich,) of 10 mM. TCEP was incubated for 30 minutes at 37 °C and 650 rpm. To prevent the reorganization of the reactive thiol groups after reduction, alkylation was performed with the electrophilic reagent iodoacetamide (IAA, Sigma-Aldrich) with a final concentration of 30 mM was also carried out for 30 minutes at 22°C and protected from light to prevent photodegradation of the IAA. The magnetic beads used (Sera-Mag SpeedBeads Carboxylate-Modified (GE Life Sciences), hydrophilic solids 50 µg/µL, and hydrophobic solids, 50 µg/µL were combined in a ratio of 1:1, first purified in ultrapure water and reconstituted in ultrapure water at a final concentration of 5 µg/µL. 2 µL of the purified bead solution was added to 1 µg of protein. The number of beads added always corresponds to a ratio of 2 µL beads to 1 µg protein and was adjusted accordingly depending on the area cut out with the LMD. The binding of the proteins to the surface of the beads was induced by adding 40 µL of absolute ethanol, resulting in a final ethanol concentration of 50%. To complete the binding process, the suspension was incubated for 10 minutes at 24 °C and 1000 rpm. The suspension was then incubated for 2 minutes on a magnetic rack, which ensures the pelleting of the magnetic beads. The supernatant was then carefully removed, and the pellet was washed outside the magnetic rack with 60 µL of 80% ethanol. The washing step was performed a total of 5 times. After the last washing step, the supernatant was removed again, and digestion was continued.

### Digestion

For the purpose of facilitating the process of digestion, a trypsin stock solution (Promega, sequencing grade) was meticulously prepared in 50 mM ABC and added in a ratio of 1:20 (enzyme:substrate). This was undertaken to ensure optimal enzyme activity, with the total volume of digestion being 40 µL. The tubes were then subjected to an ultrasonic bath for a duration of 30 seconds and subsequently incubated for a period of 18 hours at a temperature of 37 °C and a rotational velocity of 1000 rpm. Following the incubation stage, the samples were subjected to a second incubation in the magnetic rack for a period of two minutes with the objective of pelleting the beads. Thereafter, the remaining fluid was transferred to a fresh Protein LoBind tube. After this, the pellet was washed with 40 µL ABC by means of pipette mixing to elute any remaining peptides. This was followed by centrifugation at 20,000 x g for one minute and renewed incubation on the magnetic rack for two minutes. The superior portion was extracted and amalgamated with the initial eluate. The process of enzymatic digestion was terminated by the addition of 10% trifluoroacetic acid (TFA), and the pH level was manually assessed using a pH strip, ensuring it remained below the critical pH of 3. The purified peptide solution was then subjected to vacuum drying, after which it was reconstituted in 0.1% TFA.

### Protein Analysis-Digestion Control

Prior to the analysis of the purified peptide solution by mass spectrometry, a volume of 2.5 µL from each sample was injected onto a monolithic liquid chromatography column (Thermo Fisher™ PepSwift Monolithic RSLC; 200 µm x 5 cm) with an upstream trap column (Thermo Fisher™ PepSwift Monolithic Trap; 200 µm x 5 mm) coupled with UV detection to verify complete digestion.

### LC-MS/MS analysis of protein samples

A volume of 2.5 µL of each sample was loaded onto a Vanquish Neo™ UHPLC system (Thermo Fisher), which was coupled with an Orbitrap Astral™ mass spectrometer (Thermo Fisher), operated in data-independent acquisition (DIA) mode. The separation of the peptides was achieved by means of an Ionopticks™ Aurora Ultimate™ 25×75 XT C18 UHPLC column. The separation process was conducted within an active gradient of 30 minutes, with a flow rate of 500 nL/min for the initial 2 minutes and 250 nL/min for the subsequent 28 minutes. The gradient was programmed as follows: 1.0% B – 4.0% B in 0.1 minutes (500 nL/min), 4.0% B – 8.0% B in 1.9 minutes (500 nL/min), 8.0% B – 30.0% B in 25 minutes (250 nL/min), and 30.0% B – 44.0% B in 3 minutes (250 nL/min). MS data acquisition was performed in DIA (data-independent acquisition) mode. The full scan (MS1) spectra were recorded within a mass range of 400–800 *m/z* with a resolution of 240,000 @ 200 *m/z*. The automatic gain control (AGC) target was set to 5E6 for the MS1 scan, with a maximum injection time of 100 milliseconds. For the DIA analysis, the precursor mass range between 400–800 *m/z* was selected, with an isolation window of 2 *m/z*. During the scan process, the AGC target was set to 8E4 with a maximum injection time of 3 milliseconds. The fragmentation process was executed by means of higher-energy collision dissociation (HCD), with the normalized collision energy set to 25% (NCE).

### Protein data processing

For the analysis of the spectra collected in DIA mode, the data was uploaded to the Spectronaut (Biognosys) software and evaluated using a direct-DIA (library-free) approach.For the analysis of the samples, the data was introduced to the Spectronaut software (Biognosys) and analyzed with a direct DIA-based search. As protein database, the UniProt mouse reviewed database, accessed on 28^th^ August 2023 containing 17,175 entries, was selected. Search and extraction settings were kept as standard (BGS Factory settings) Normalization was done by the software, using global normalization based on the median. For reliable label-free quantification, only proteins identified with ≥ 2 unique peptides were considered for further analysis. Subsequently, the average normalized abundances (determined using Spectronaut) were calculated for each protein and used to determine the ratio between the metastatic tissue and non-affected tissue. Only proteins with a p-value of ≤0.05 and an abundance ratio of ≥2 or ≥0.5 (up-regulated as well as down-regulated) were considered as finally regulated.

Functional enrichment analysis was performed using the STRING database (version 12.0; https://string-db.org/). Kyoto Encyclopedia of Genes and Genomes (KEGG) pathway enrichment was conducted based on the integrated KEGG database resource within STRING. Identified pathways were ranked according to the STRING *strength* score, which reflects the enrichment magnitude relative to expected background. Default parameters were applied. Venn diagrams were created using a web-based tool^35^.

## Acknowledgements

A.F., A.C.G. and K.W.S. acknowledges funding from Federal Ministry of Research, Technology and Space (Bundesministerium für Forschung, Technologie und Raumfahrt; BMFTR) under the funding reference 161L0271. A.S. and S.H. acknowledge funding from Federal Ministry of Research, Technology and Space (Bundesministerium für Forschung, Technologie und Raumfahrt; BMFTR) and support by the Ministry of Culture and Science of the State of North Rhine-Westphalia (Ministerium für Kultur and Wissenschaft des Landes Nordrhein-Westfalen, MKW NRW). A.T. acknowledges the support of an Emmy Noether Award from the German Research Foundation (DFG, 467788900) and the Ministry of Culture and Science of the State of North Rhine-Westphalia (NRW-Nachwuchsgruppenprogramm). A.T. acknowledges the support of an ERC starting grant (METATARGET, 101078355). A.T. holds the Peter Hans Hofschneider endowed Professorship of Molecular Medicine from the Stiftung Experimentelle Biomedizin. The authors would like to thank Yanina Dening for her laboratory expertise.

## Author Contributions

K.W.S., A.S., S.H. and A.T. conceived the study. K.H., A.F., F.L.H., A.C.G., A.H, J.C. and K.W.S. generated and analyzed all data. L.M.N.M., F.C. and G.A. generated and provided sample material. K.H., A.F., A.H., S.H. and K.W.S. wrote the original manuscript. All authors contributed to editing and revision of the final manuscript. K.W.S., A.S., S.H. and A.T. supervised the work.

