## Supplementary figures for "SIMO – Single Section Integrative Multi-Omics – spatial mapping of metabolites and lipids combined with region-specific proteomics in a single tissue slice"

Figure S1 Hau and Fecke at al.

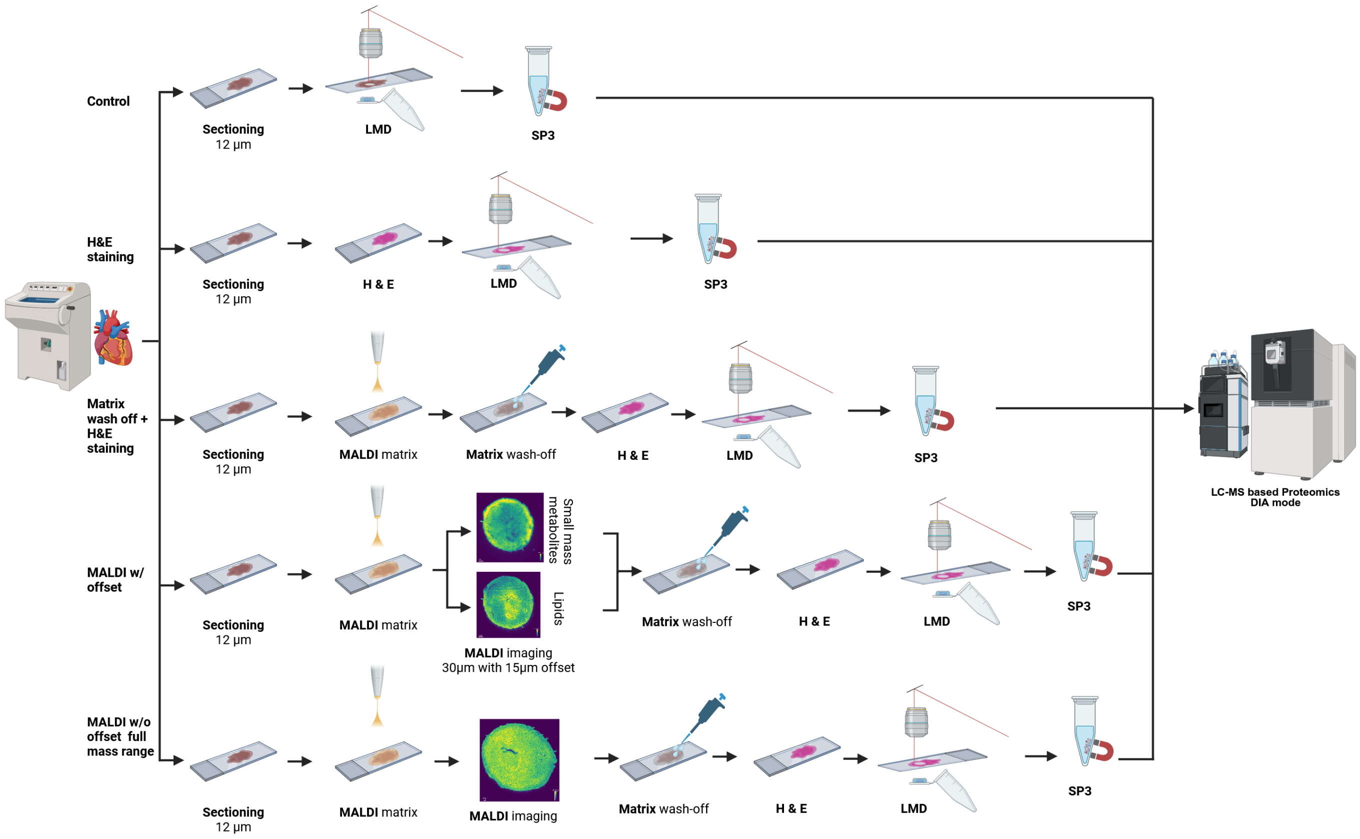


Supplementary Figure S1: Schematic overview of the stepwise method development workflow using heart samples. Individual processing steps (LMD excision, H&E staining, matrix removal and MALDI imaging) were successively added and evaluated by comparing protein identification numbers from LC-MS-based proteomics analysis.

Figure S2 Hau and Fecke at al.

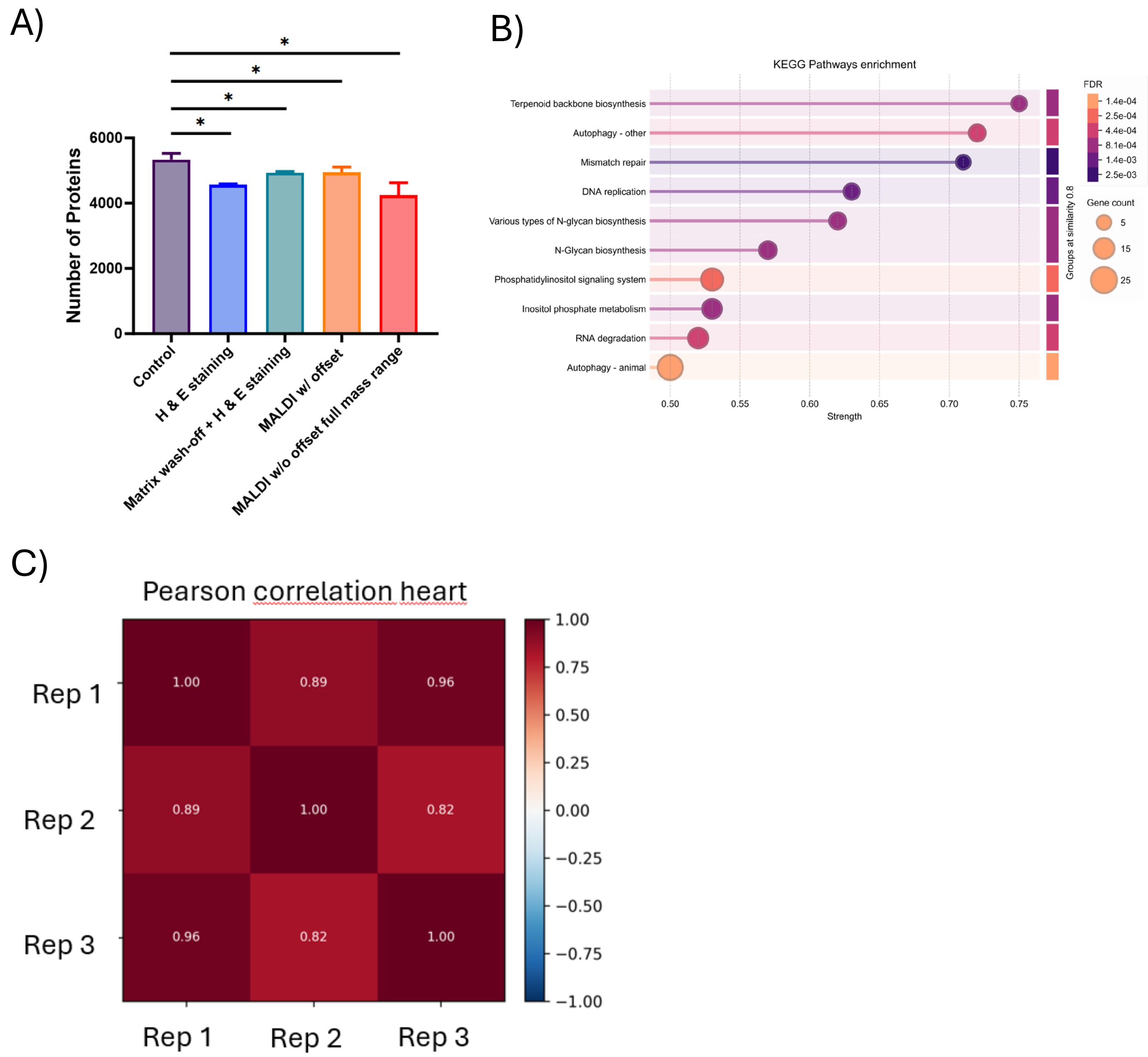


Supplementary Figure S2 A) The bar plot shows the average number of identified proteins across the individual method development steps with their standard deviation. Control samples of all method development steps were pooled together (n = 9). ). Ordinary one-way ANOVA was performed against control tissue to determine statistic significance (*p < 0.05). B) KEGG Pathways enrichment analysis using STRING database (Version 12.0) based on the input protein set (1211 proteins) of identified proteins only present in control tissue. Pathways are ranked according to the STRING strength score. The displayed pathway showed significant enrichment (FDR ≤ 0.05). C) Pearson correlation of triplicate MALDI-MSI heart data

Figure S3 Hau and Fecke at al.

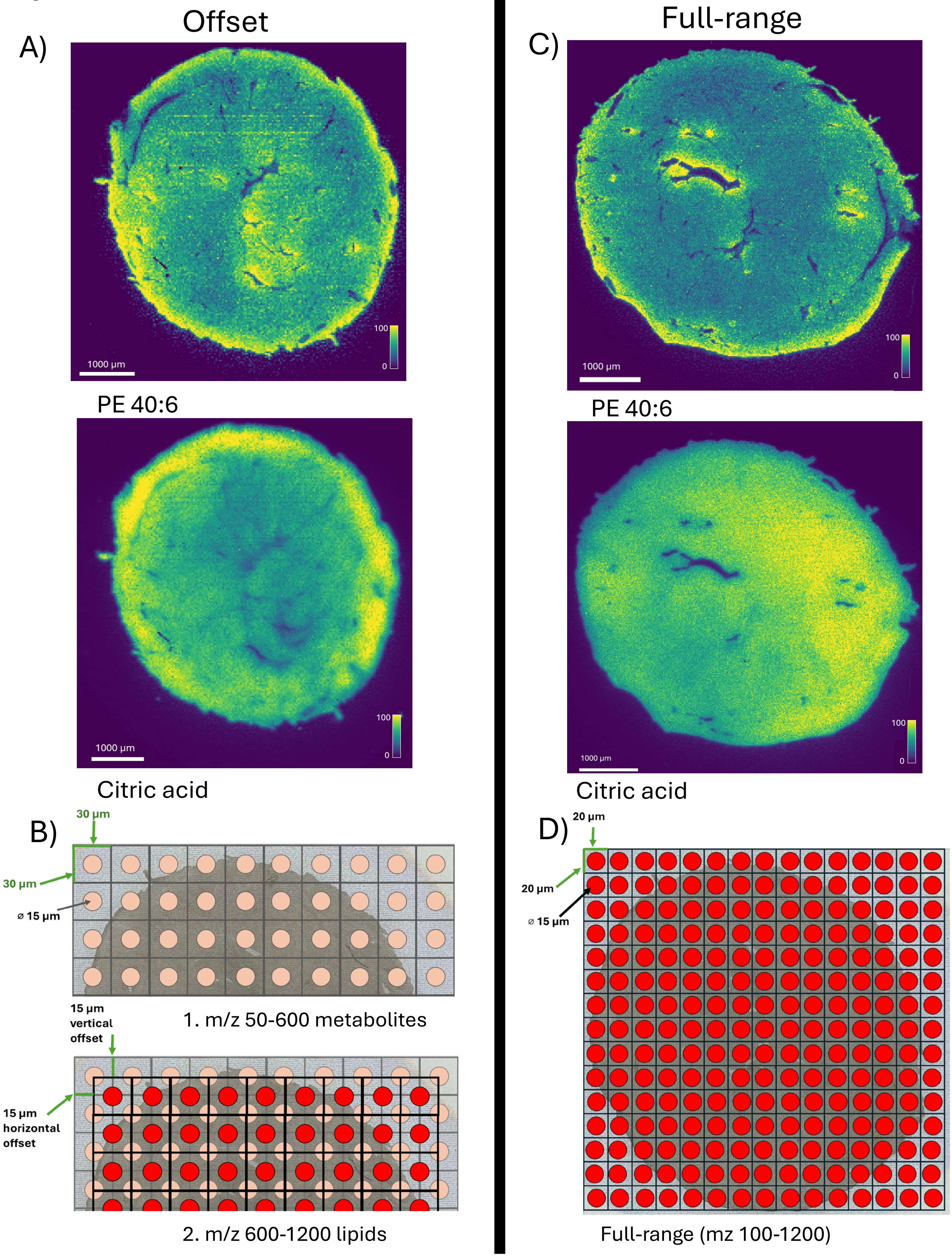


Supplementary Figure S3 A) Exemplary MALDI MS images of metabolites and lipids measured in consecutive measurements with a laser-offset in heart tissue. Signal intensities are indicated by the color scale B) schematic representation of offset MALDI measurements. First the metabolite measurement was performed from m/z 50-600 at 30x30 µm pixel size. Next, lipids were measured by re-measuring the same sample with a 15x15 µm laser offset from m/z 600-1200. C) Exemplary MALDI MS images of metabolites and lipids measured simultaneously in a single measurement in heart tissue. Signal intensities are indicated by the color scale. D) Schematic representation of full-range MALDI measurements. Metabolites and lipids are captured simultaneously from m/z 100-1200 at 20x20 µm spatial resolution.

Figure S4 Hau and Fecke at al.

### Heart post LMD

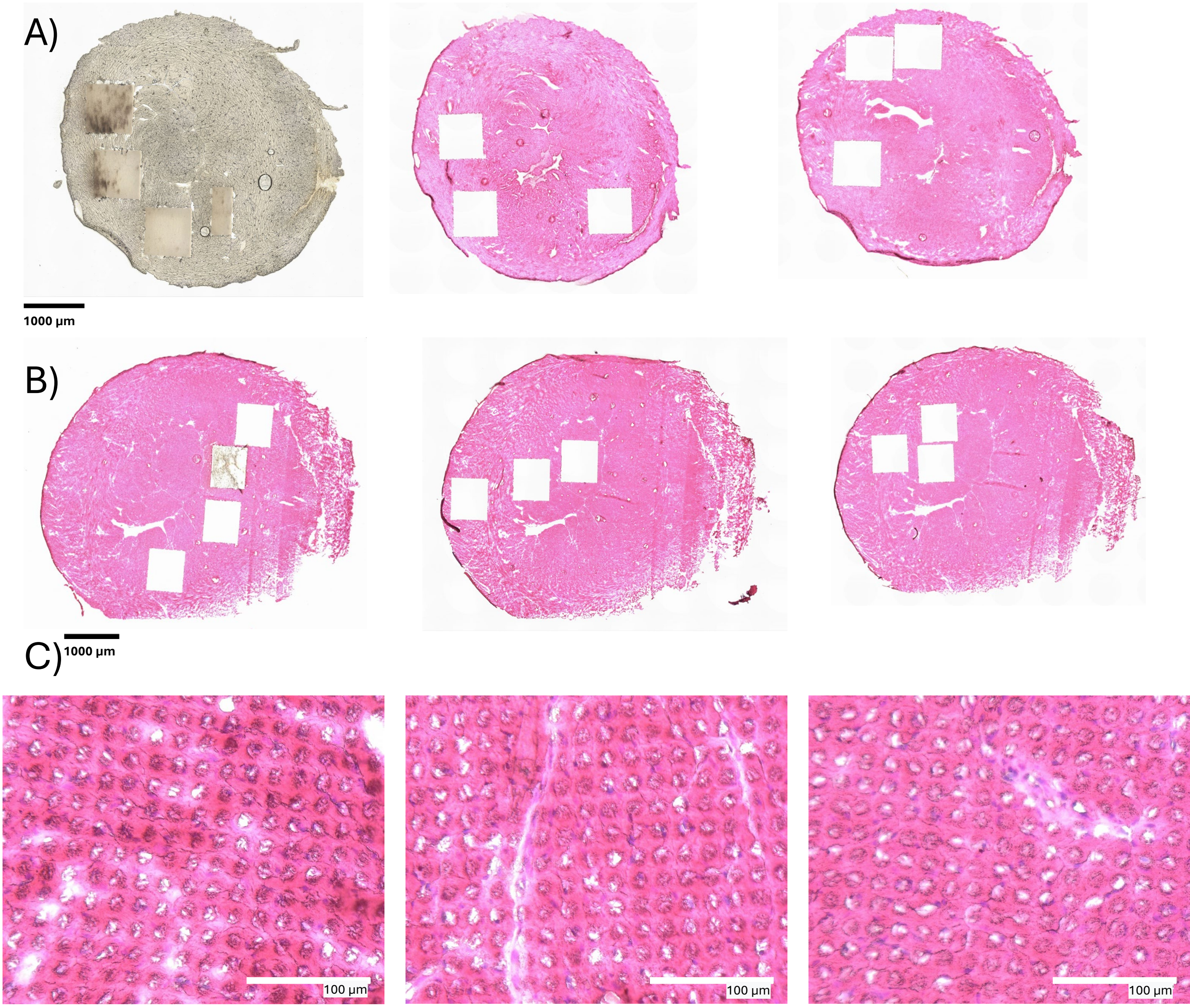


Supplementary Figure S4: Post-LMD Microscopy of heart sections used for method optimization. A) shows an unstained control heart section, a stained heart section measured with offset and a stained heart section measured at full range, respectively. B) shows final stained triplicate heart sections measured for method validation. C) Zoom-in into H&E stained triplicate heart sections post MALDI.

Figure S5 Hau and Fecke at al.

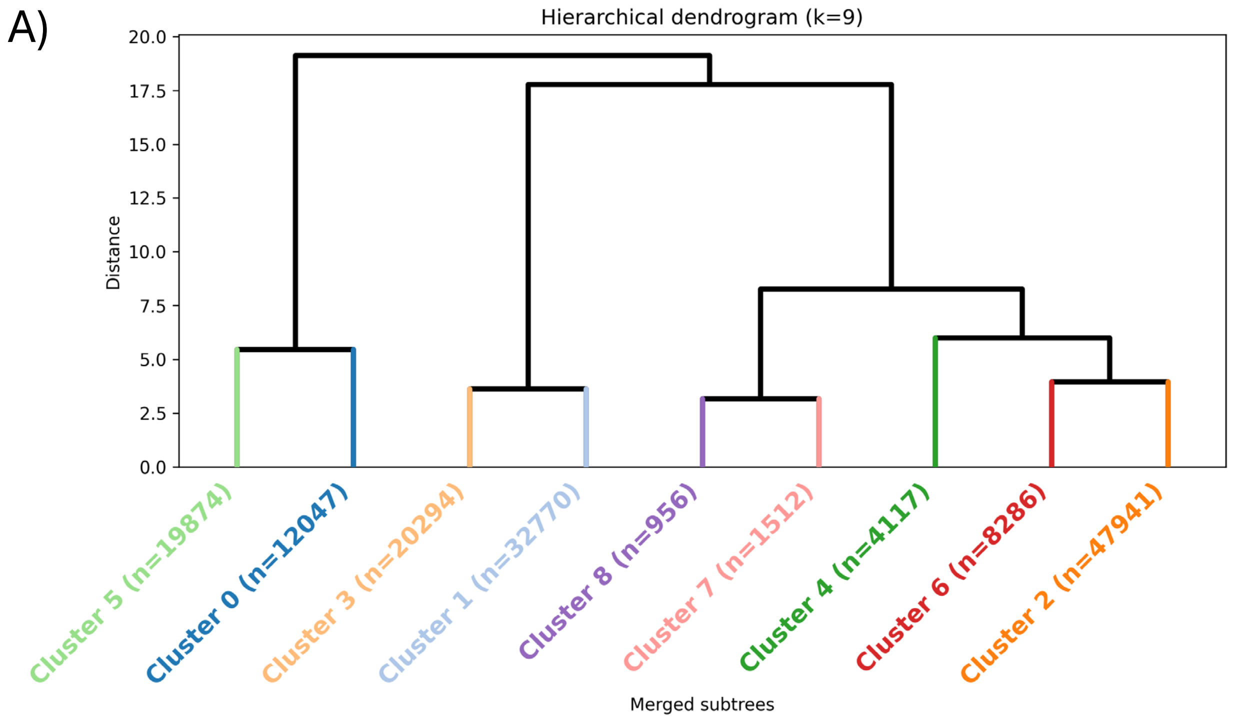


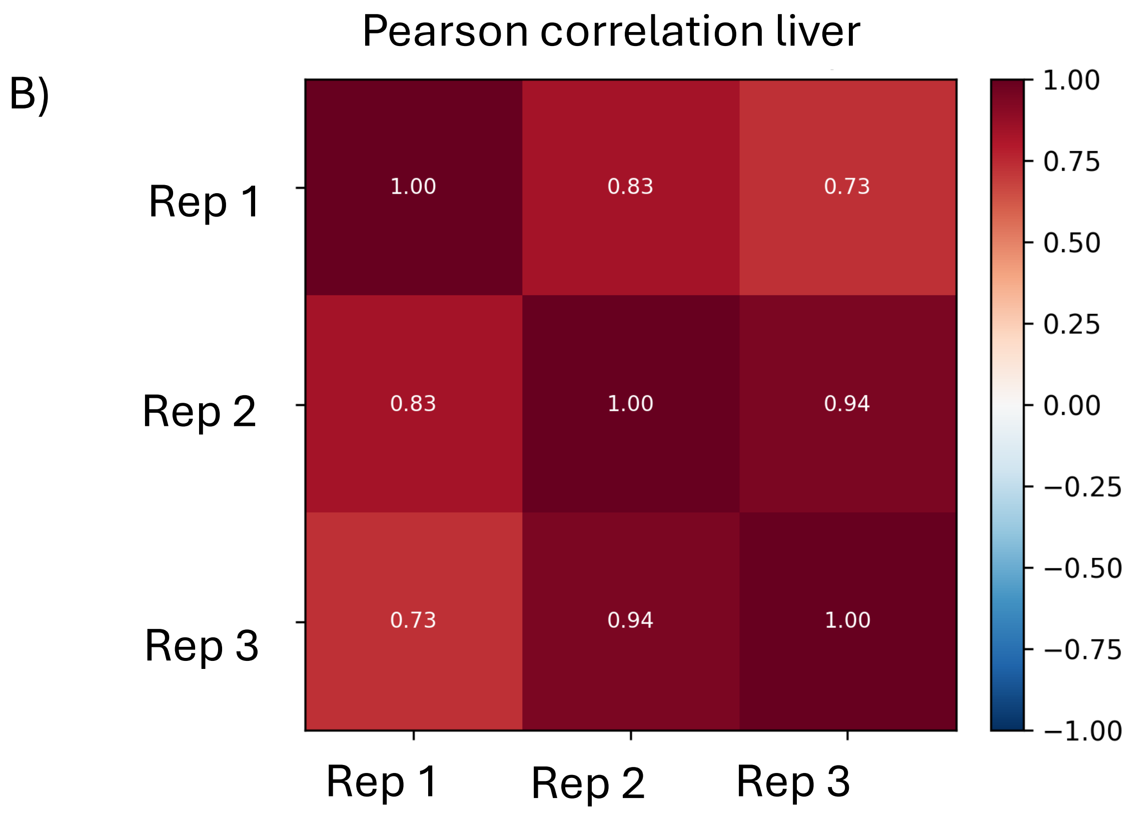


Supplementary Figure S5:A) Dendrogram illustrating the results of hierarchical clustering of liver tissue with nine clusters. Pixels are grouped by molecular similarity, forming clusters by stepwise merging to minimizing variance. Each leaf represents a merged pixel group colored by cluster, and branch height indicates the dissimilarity between clusters B) Pearson correlation of MALDI-MSI data in liver replicates

Figure S6 Hau and Fecke at al.

#### A) Metabolites with higher abundance in tumor

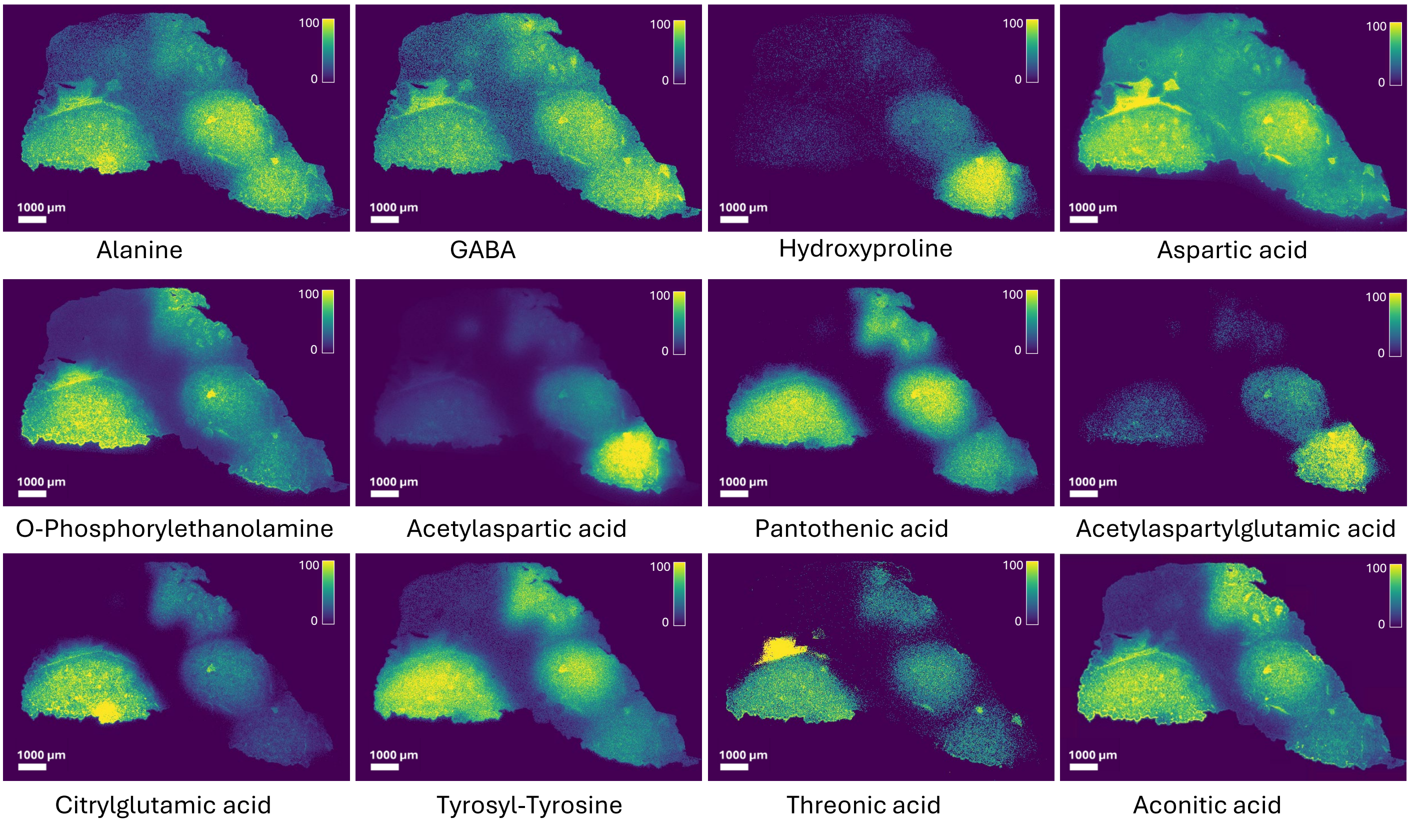


#### B) Metabolites with lower abundance in tumor

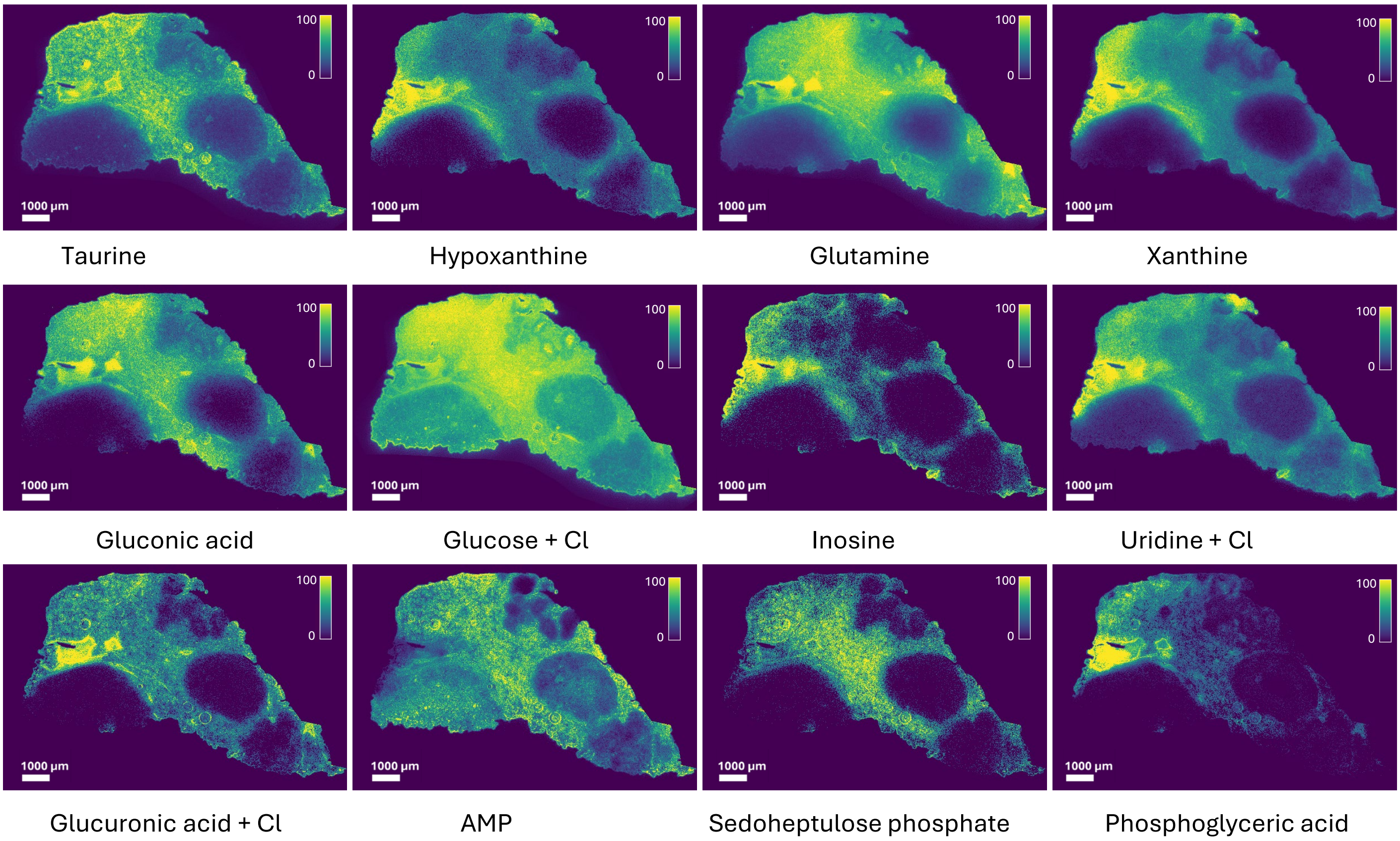


Supplementary Figure S6: Exemplary MALDI MS images of metabolites with A) higher abundance in tumor regions and B) lower abundance in tumor regions. Signal intensities are indicated by the color scale

Figure S7 Hau and Fecke at al.

#### A) Lipids with higher abundance in tumor

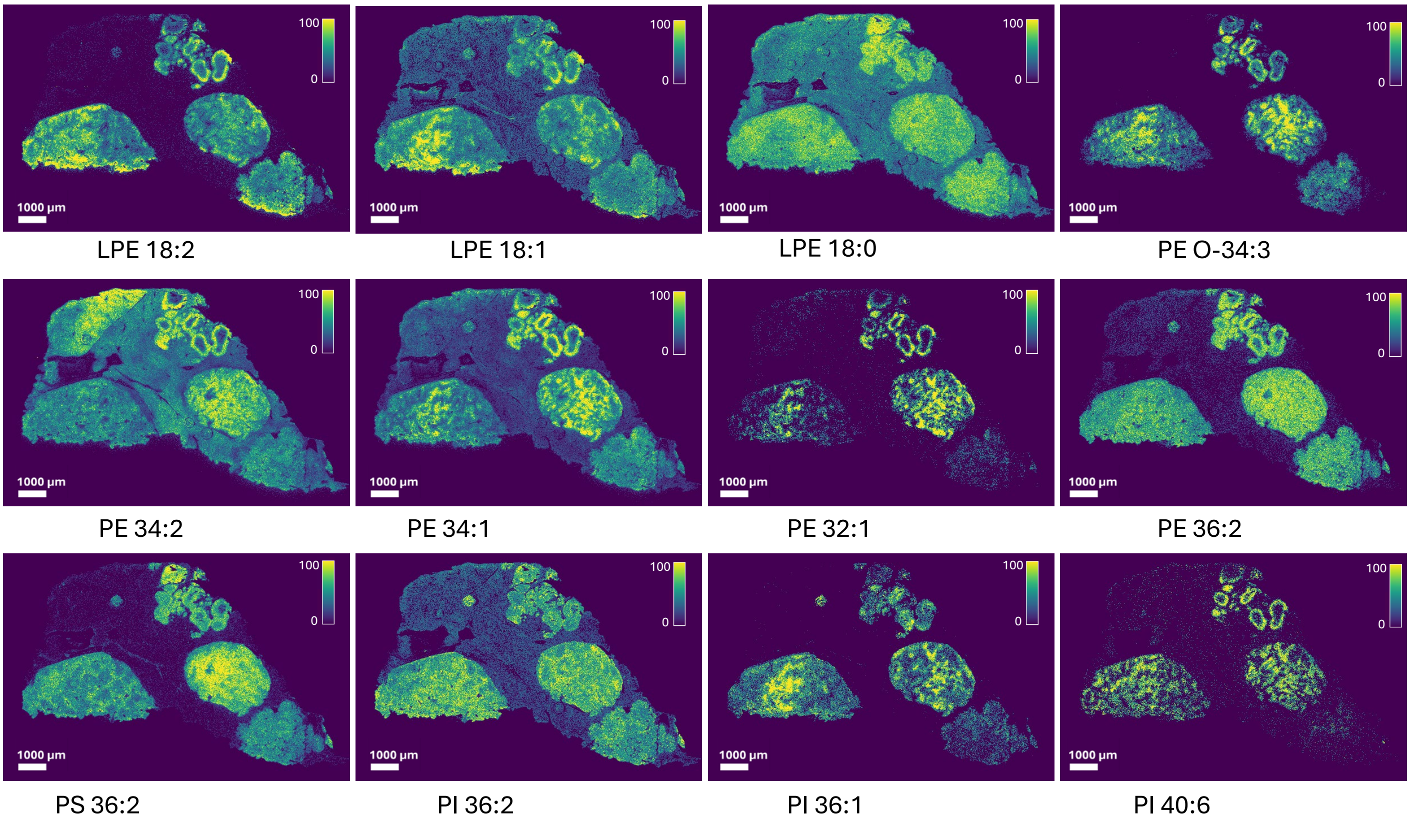


#### B) Lipids with lower abundance in tumor

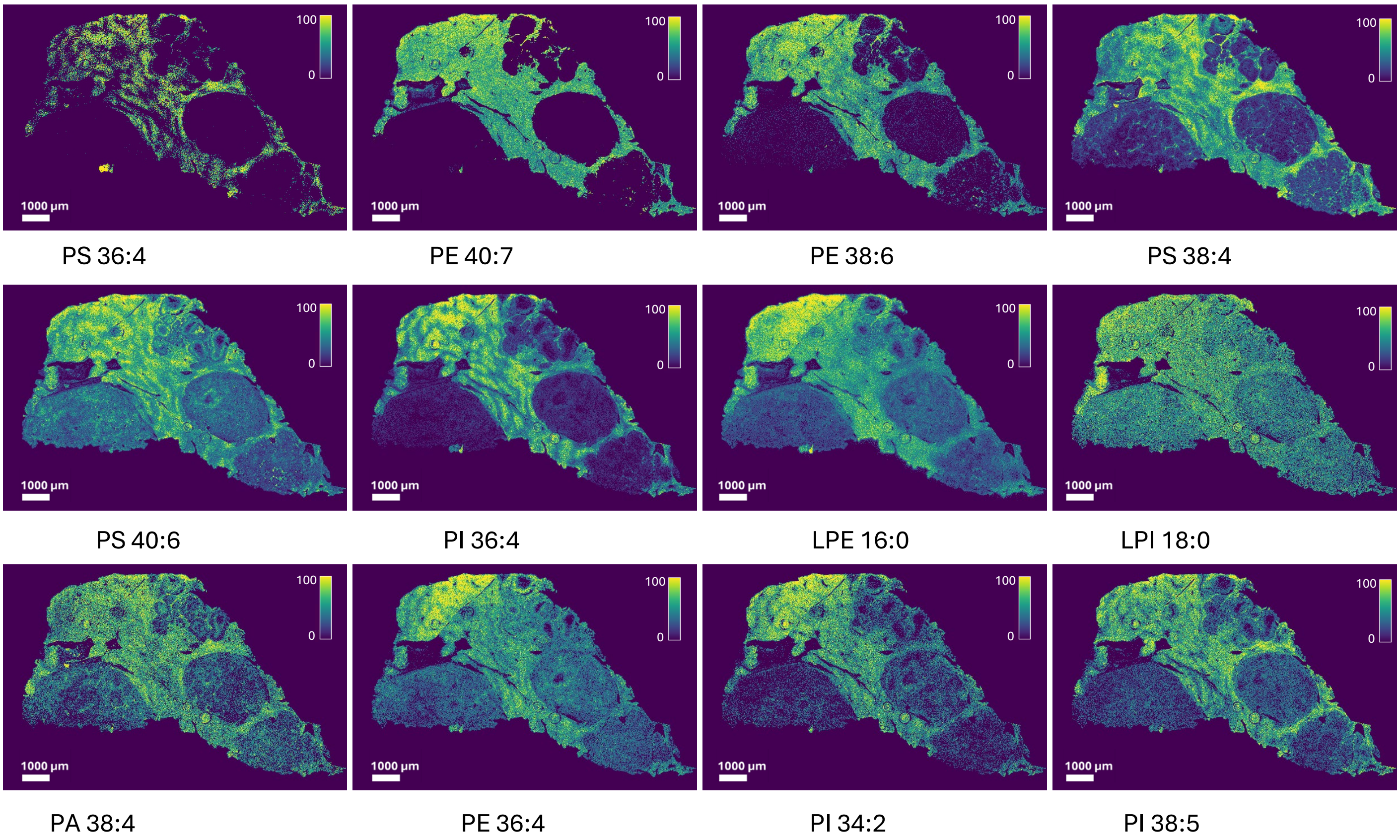


Supplementary Figure S7: Exemplary MALDI MS images of lipids with A) higher abundance in tumor regions and B) lower abundance in tumor regions. Signal intensities are indicated by the color scale

Figure S8 Hau and Fecke at al.

A)

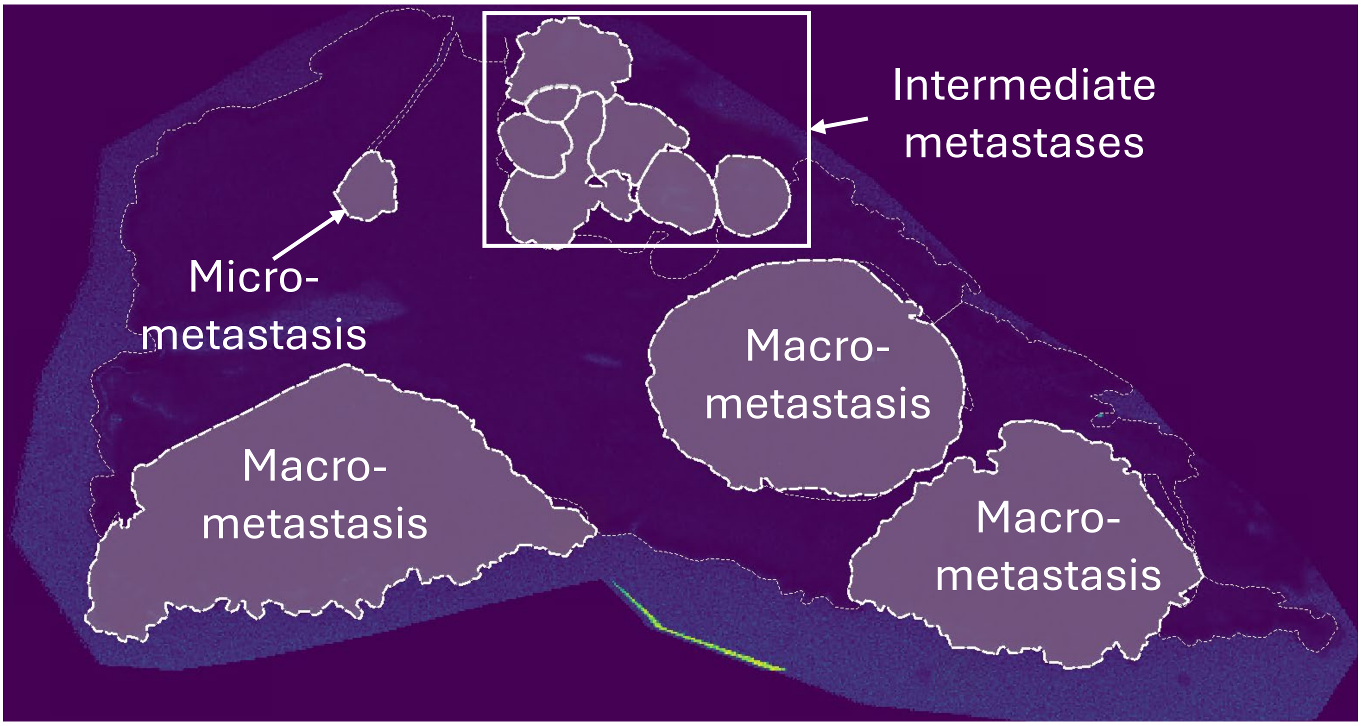


B)

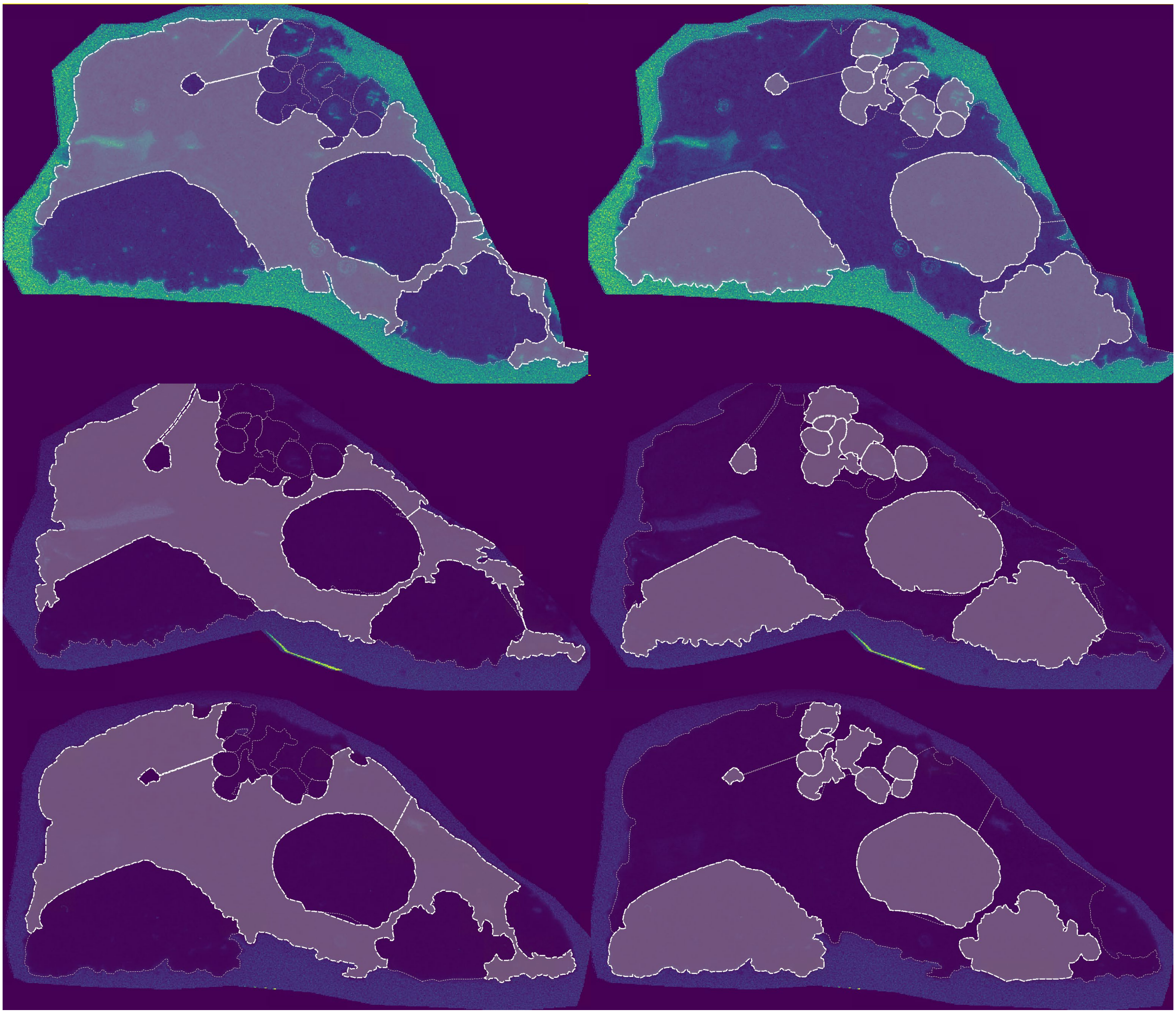

Supplementary Figure S8: MALDI-ROI analysis A) MALDI-ROI grouping for statistical analysis B) Manually drawn ROIs in replicate metastatic liver sections showing adjacent non-tumor tissue (left) and drawn tumor regions (right).

Figure S9 Hau and Fecke at al.

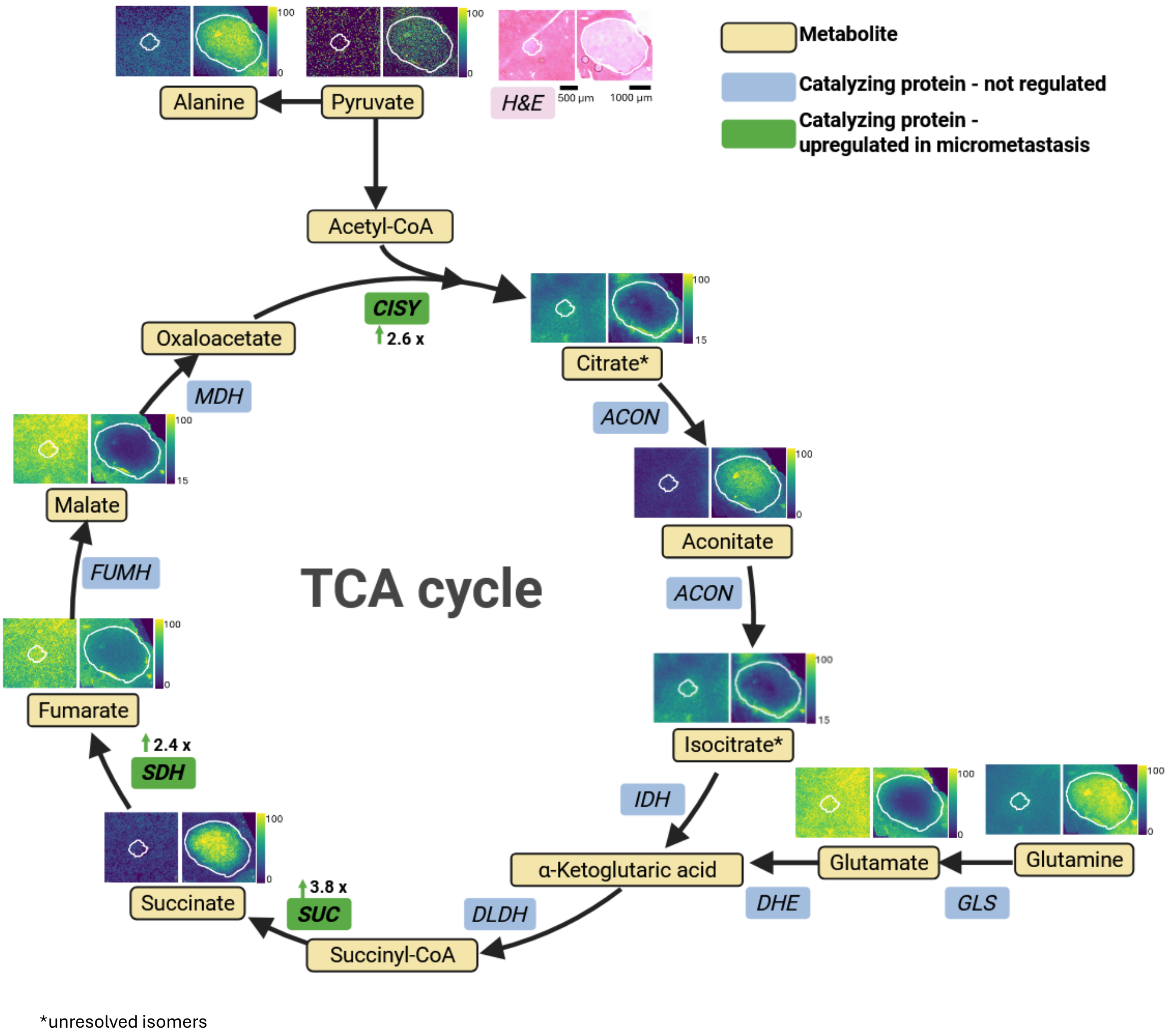


Supplementary Figure S9: Schmatic representation of TCA-cycle and aminoacid metabolite spatial distributions obtained from MALDI-MSI in micrometastasis 1 and macrometastasis 6. Signal intensities are indicated by the colour scale. Unresolved isomers are highlighted with *. Corresponding catalyzing proteins are shown alongside to demonstrate pathway activity. Proteins displayed in green idicate upregulation in micrometastasis compared to macrometastasis. Post-MALDI H&E shows histological tumor regions.

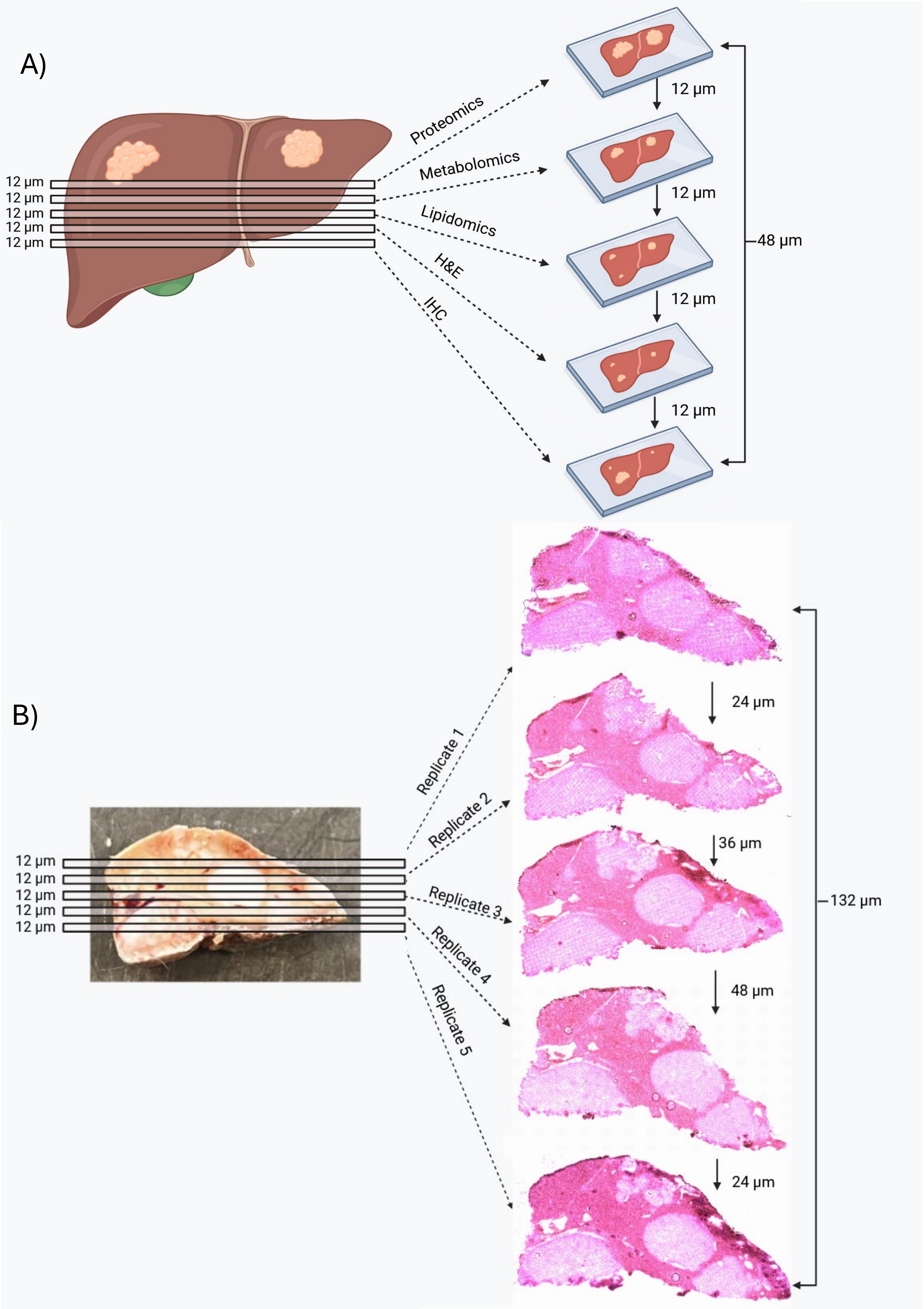


Supplementary Figure S10: A) Schematic representation of sample requirements for serial-section-based Multi-omics workflows. The sampled tissue area and cell layer changes with every serial section, reducing comparability of the different analyses carried out. B) Representation of performed serial sectioning for replicate MALDI-MSI analysis.

Figure S11 Hau and Fecke at al.

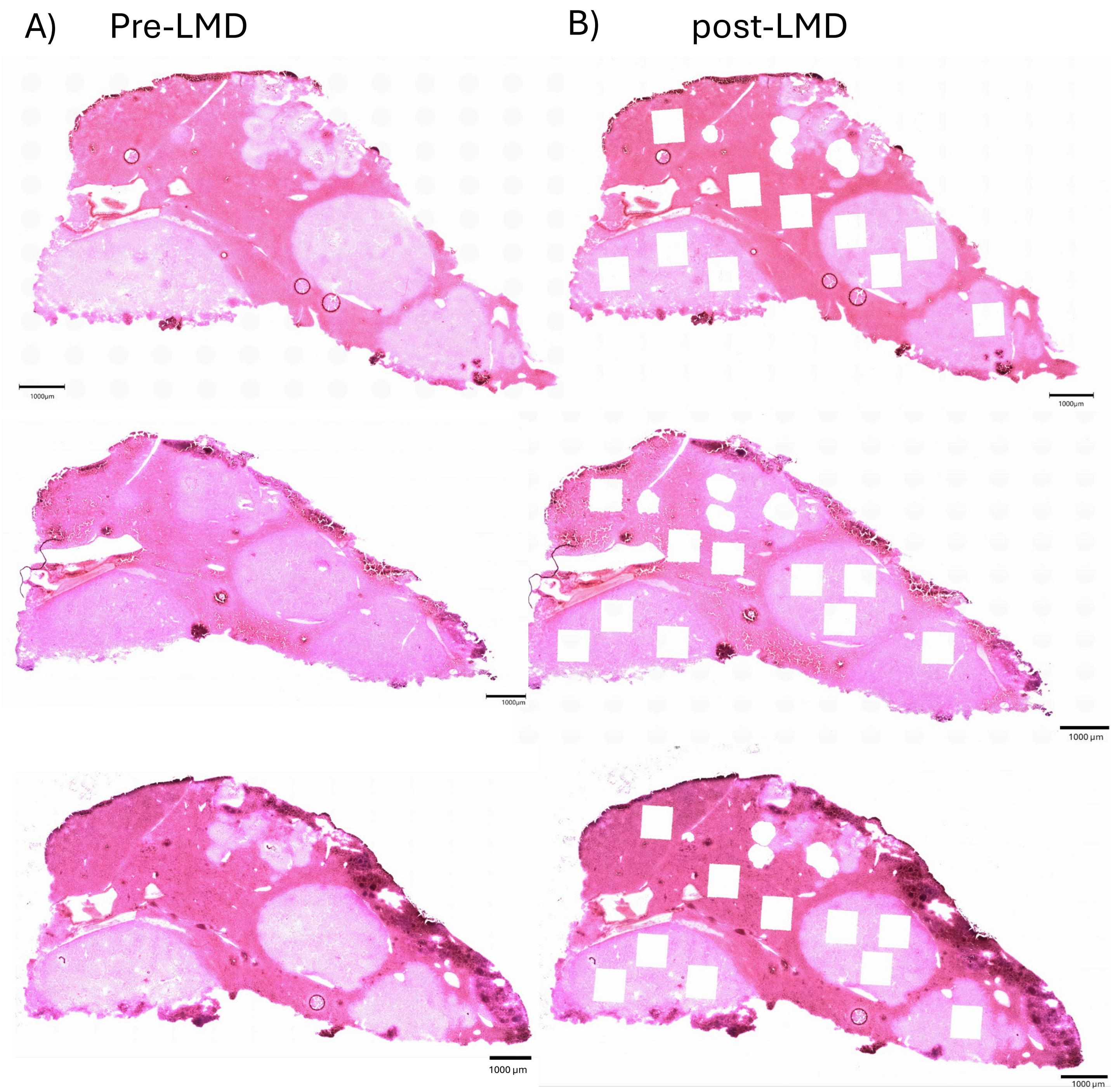


Supplementary Figure S11: A) H&E-stained sections of metastatic liver triplicate sections B) H&E-stained sections of metastatic liver sections post LMD, depicting excised areas.
